# Metabolic Salvage and Acyl-chain Remodeling Support Glycosphingolipid Synthesis within the PDAC Tumor Microenvironment

**DOI:** 10.64898/2026.04.14.718544

**Authors:** Anna S. Trimble, Casie S. Kubota, Elaine Zhao, Maureen L. Ruchhoeft, Jonathan R. Weitz, Woncheol Jung, Kristina L. Peck, Satoshi Ogawa, Ethan L. Ashley, Herve Tiriac, Tae Gyu Oh, Andrew M. Lowy, Dannielle D. Engle, Christian M. Metallo

**Author notes:** Correspondence: Dannielle Engle, Christian Metallo.

## Abstract

Pancreatic ductal adenocarcinoma (PDAC) is a highly lethal malignancy where metabolic homeostasis is maintained by tumor and stromal cells within the tumor microenvironment (TME). To better assess pathways supporting macromolecule biosynthesis in PDAC tumors, we apply ^13^C metabolic flux analysis (MFA) to slice cultures of treatment-naïve human tumors and mouse models that retain the native TME. Glycans, lipid headgroups, and very long-chain fatty acids are the most dynamic metabolic pools, while long chain fatty acids, purines, and pyrimidines are predominantly salvaged locally *in situ*. We use targeted pharmacological modulators to highlight the importance of recycling pathways and metabolic redundancies which mitigate changes in lipid abundances. Finally, we leverage targeted lipid fluxomics and the distinct ganglioside and globoside profiles of tumor and stromal cells, respectively, to demonstrate the role of the lipid kinase PIKfyve in supporting ganglioside homeostasis via sialic acid and ceramide salvage. These data establish application of MFA to slice cultures of PDAC tumors as an effective approach for assessing metabolic mechanisms and therapeutic responses within an intact TME.

## Introduction

Pancreatic ductal adenocarcinoma (PDAC) is a deadly malignancy notorious for its poor prognosis. The development of more targeted therapeutics requires a deeper understanding of the specific mechanisms through which PDAC tumors sustain growth and survival. Aberrant metabolism is a hallmark of cancer^1–3^, but the PDAC tumor microenvironment (TME) is characteristically nutrient sparse, with poor vascular perfusion^4–6^ and limited substrate availability^7–10^. Therefore, PDAC tumor cells must negotiate a range of metabolic stresses to survive and proliferate. Proliferation requires the synthesis of new macromolecules like nucleic acids, proteins, glycans, and lipids, which, in turn, require adequate supplies of the purine, pyrimidine, amino acid, sugar, and fatty acid precursors from which they are synthesized. When nutrients are scarce, salvage pathways become particularly important. For example, PDAC cells scavenge amino acids from extracellular protein^8,11,12^, recycle N-acetyl-glucosamine (GlcNAc) and uridine^13^, and rely on extracellular lipids and autophagy^14,15^ to maintain metabolic homeostasis in nutrient-deficient or hypoxic conditions. Sustaining macromolecule synthesis is critical for tumor progression, making these pathways an attractive therapeutic target for most cancers. However, translation of these findings to the clinic remains challenging. Treatments targeting these metabolic nodes, such as the pyrimidine analogs gemcitabine and 5-fluorouracil, show only limited efficacy in PDAC, and clinical trials targeting salvage mechanisms like autophagy with compounds like hydroxychloroquine have failed^16–18^. Even with chemotherapeutic treatment, median overall survival for patients with metastatic disease remains less than one year^19^. A mechanistic understanding of how PDAC cells orchestrate macromolecule synthesis within the constraints of the TME remains unknown.

PDAC tumors are notable for their low degree of cellularity. On average, epithelial cancer cells often comprise less than half of the tumor cellular content, with stromal cells, especially fibroblasts and immune cells, comprising the remainder^20^. The presence and functions of these stromal cells contribute directly to disease progression and correlate with prognosis^10,21–23^. Previous work has demonstrated that the stroma can provide amino acids^21,22^, nucleotides^24,25^, and lipids^26,27^ to maintain PDAC metabolic homeostasis. However, studying PDAC tumor metabolism within the native TME has proven challenging. Organotypic pancreatic tissue slices have been an effective tool for studying pancreas function^28–30^, type-1 diabetes^31^, cancer drug response^32–35^, assessing new therapeutic strategies^36–38^, and gene therapy evaluation^39^. Previous work established that these slices maintain viability, cellular diversity, and transcription for at least several days in culture^32–34,40–42^, and this method has been used to investigate ketone body^43^ and BCAA^22^ metabolism in PDAC tumors.

Here, we describe a method for performing metabolic flux studies in PDAC slice cultures to quantify the contributions of *de novo* nutrient synthesis and salvage to macromolecule synthesis. Using tumor slices from multiple murine models of PDAC and from treatment-naïve human PDAC tumors, we quantify *de novo* fatty acid, amino acid, nucleotide, and nucleotide sugar synthesis as well as their incorporation into biological macromolecules. Performing concurrent metabolic flux analysis (MFA) studies within the same tumor highlights unique aspects of *in situ* PDAC physiology. Lipid turnover is exceptionally high within the PDAC tumor slices, and we note that long-chain fatty acids, purines, and pyrimidines are predominantly salvaged in both murine and human PDAC tumors. Multiple salvage pathways contribute to the maintenance of lipid homeostasis, including the lipid kinase PIKfyve which sustains the synthesis of tumor cell-specific glycosphingolipids. Collectively, these results suggest that application of metabolic flux analysis to PDAC slice culture can facilitate elucidation of active metabolic pathways and drug mechanisms in patient tumors.

## Results

### Application of Metabolic Flux Analysis to PDAC Slice Cultures

To quantify synthesis and salvage fluxes of PDAC within the native TME, we performed stable isotope tracing in PDAC tumor slices from mice bearing autochthonous or orthotopically grafted intact piece (OGIP) KPC (Kras^G12D^, Trp53^R1725H^, Pdx1-Cre) tumors (Fig. 1A). Expanding a single KPC tumor into multiple OGIP tumors enables the investigation of additional time points and conditions while restraining the degree of intertumoral variability. Immunohistochemistry (IHC) staining for the fibroblast marker podoplanin and macrophage marker F4/80 demonstrate that all tumors had representation of these TME constituents (Fig. 1B-D, Extended Data Fig. 1A), as expected^44^. Proliferation of the primary tumors was maintained at similar rates in slice culture (Fig. 1E,F). Overall, these data validate the suitability of these slice cultures for metabolic analyses through the comparability of autochthonous and OGIP KPC tumor slices and their continued proliferation in *ex vivo* culture.

**Fig 1.**
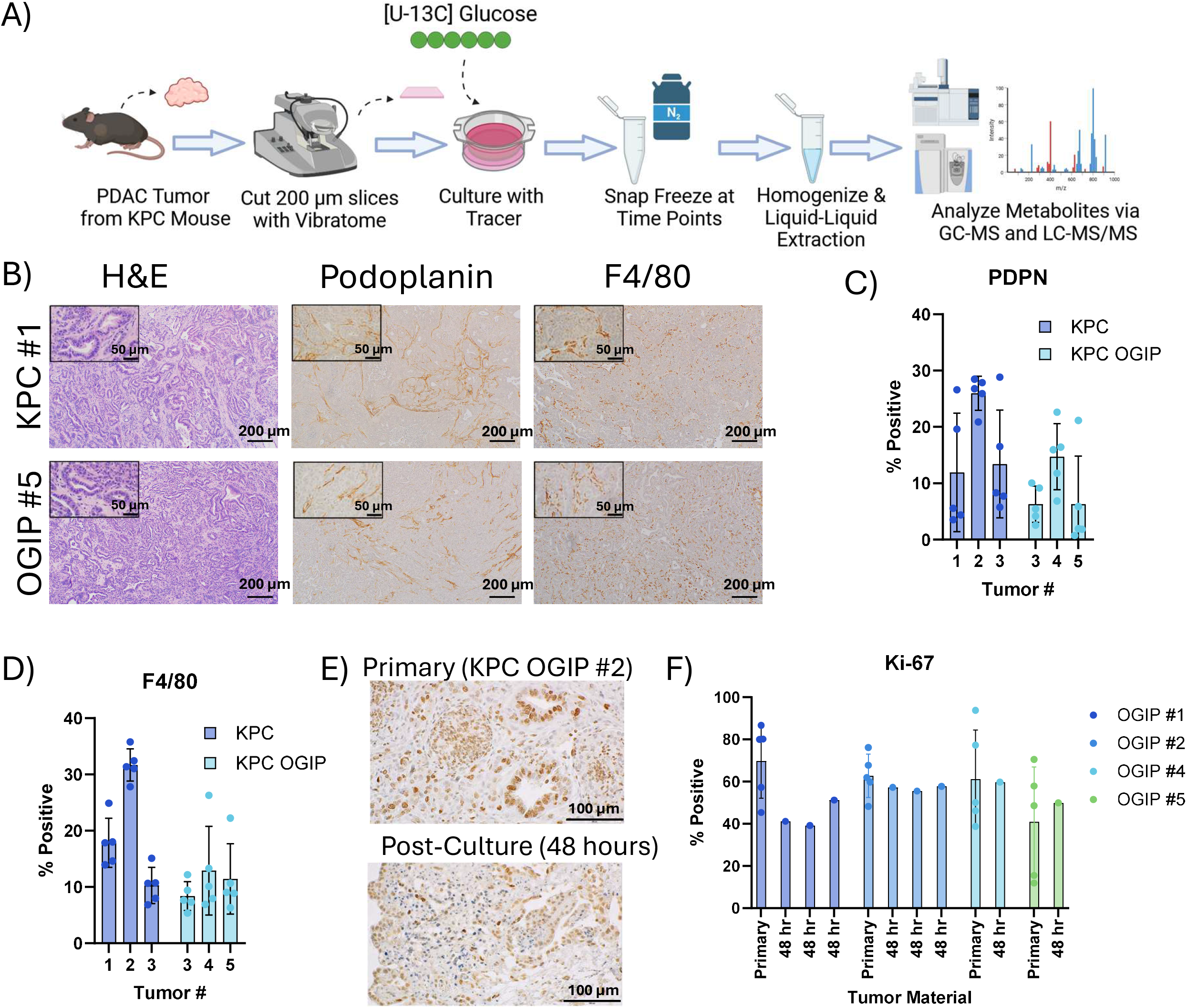
*Ex vivo* tissue slices from KPC and KPC OGIP tumors are suitable for metabolic flux studies. (A) Summary of workflow for conducting metabolic flux studies using stable isotope tracing in *ex vivo* PDAC tumor slice cultures. (B) Immunohistochemistry images for primary tumor material from a representative KPC tumor and a representative KPC orthotopically grafted in pancreas (OGIP) tumor stained for hematoxylin and eosin (H&E), podoplanin, or F4/80. Scale bars represent 200 μm. Inset scale bars are 50 μm. (C) Quantitation of the percent of cells positive for podoplanin (PDPN) or (D) F4/80 in primary tumor material. OGIP #1 and #2 were not analyzed. Data points represent independent fields of view (n=5). (E) Immunohistochemistry images showing Ki-67 in primary KPC OGIP tumor and tissue slice after 48 hours of *ex vivo* culture. Scale bars represent 100 μm. (F) Quantitation of Ki-67 in primary tumor material and post-culture tissue slices. OGIP #3 was not analyzed. For primary tumor material, data points represent independent fields of view (n=5). For post-culture material, entire slice area was quantified. All error bars show standard deviation.

The synthesis of proteins, nucleic acids, glycans, and lipids is fundamentally intertwined with the metabolism of the monomers from which they are derived. Thus, quantifying the synthesis of these macromolecules and their building blocks requires the analysis of multiple metabolite pools, ideally from the same tumor. As different metabolites require distinct analytical procedures and clinical material is limited, we designed an extraction workflow to facilitate comprehensive MS profiling from a single tumor slice (Extended Data Fig. 1B). Slice homogenate is split into three different analytical workflows: a modified Bligh and Dyer extraction for polar metabolites, biomass, and saponified fatty acids^45^, protein quantitation by BCA assay, and complex lipid analysis on LC-MS/MS. Using this approach, the labeling of precursor pools and intact macromolecules can be quantified simultaneously to capture a more complete picture of *in situ* metabolism within each slice.

To obtain a baseline metabolic profile of PDAC within the TME, we cultured tumor slices from autochthonous or OGIP KPC tumors in medium containing 25 mM [U-^13^C] Glucose (U-Glc) for 24, 48, or 96 hours. These slices were metabolically active, as we observed high levels of TCA cycle intermediate labeling within 24 hours (Fig. 2A). As glucose is also a key carbon source for the synthesis of non-essential amino acids (NEAAs) (Fig. 2B), we examined amino acid labeling. Alanine was highly labeled from U-Glc, with nearly 75% of the free alanine pool enriched at 24 hours (Fig. 2C). Additionally, nearly 30% of the aspartate pool and around 40% of the glutamate pool were enriched by U-Glc within 24 hours. α-ketoglutarate (aKG) and malate were similarly labeled, suggesting these amino acids are highly synthesized in PDAC slices. Other NEAAs, including serine, glycine, asparagine, and proline, were also labeled from U-Glc, albeit at lower levels that remained relatively constant over time. Importantly, both TCA and NEAA labeling were consistent across multiple autochthonous and OGIP KPC tumors (Extended Data Fig. 2A-D).

**Fig 2.**
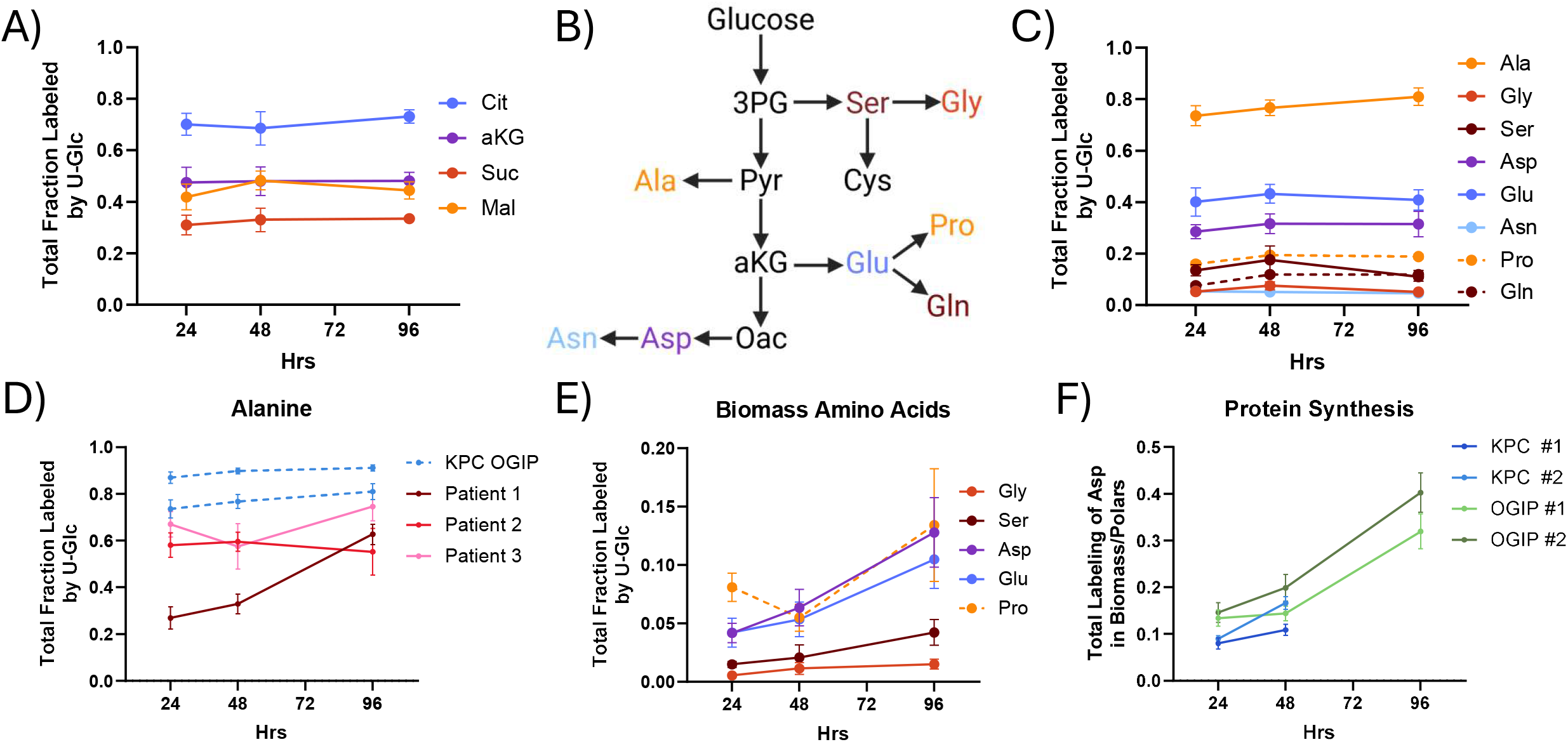
Quantifying Protein Synthesis in KPC Tissue Slices. (A) Total fraction labeled of TCA cycle intermediates from U-Glc in tumor slices from a representative KPC OGIP tumor (OGIP #2). (B) Diagram summarizing the *de novo* synthesis of non-essential amino acids from glucose. (C) Total fraction labeled by U-Glc for non-essential amino acids in KPC OGIP #2 tumor slices. (D) Total labeling of alanine from U-Glc in human PDAC slices. KPC OGIP #1 and #2 shown for reference. (E) Total fraction labeled by U-Glc of non-essential amino acids from U-Glc in the hydrolyzed biomass of tissue slices from OGIP #2. (F) Protein synthesis over time in KPC tumor slices calculated as the total enrichment of the aspartate in the hydrolyzed biomass normalized to the total enrichment of the polar aspartate pool.

We assessed whether our findings in KPC tumors were relevant to human disease by generating tissue slices from surgically resected tumor material from three treatment-naïve human patients (Extended Data Fig. 2E). Alanine, aspartate, and glutamate enrichment were high relative to other amino acids in human PDAC slice as well (Fig. 2D). Interestingly, human PDAC slices had lower total alanine labeling than KPC tumor slices (Extended Data Fig. 2F). While the alanine labeling in most KPC tumors was greater than 80% at 48 hours, alanine labeling remained less than 60% in human PDAC slices. As stromal cells have been shown to influence alanine metabolism^21,46^, how the murine and human TME contributes to this difference remains to be explored.

Next, we quantified the protein synthesis rates via the incorporation of labeled amino acids into biomass. Proteins remaining after slice extraction were hydrolyzed to individual amino acids, and the labeling of each amino acid from U-Glc was quantified by GC-MS (Extended Data Fig. 2G). Importantly, the labeling of proteogenic NEAAs increased over time in *ex vivo* culture (Fig. 2E, Extended Data Fig. 2H), consistent with the continuing viability of these slice cultures. As the free alanine and aspartate pools were particularly enriched, the incorporation of these labeled amino acids into biomass presents a convenient method for quantifying protein synthesis. When normalized to the labeling of the respective free pool, we estimate that around 10% of the protein in slices is newly synthesized in these slice cultures by 48 hours (Fig. 2F, Extended Data Fig. 2J). Overall, this establishes that PDAC slices synthesize amino acids and protein in *ex vivo* culture.

### Nucleobase salvage supports nucleic acid and glycan synthesis

Nucleic acid synthesis is critical for tumor growth and is a common target of chemotherapeutic agents used to treat PDAC tumors. Pyrimidines and purines are both synthesized as nucleotides using metabolite pools that are enriched by U-Glc (Fig. 3A). Purine nucleotides were highly labeled by U-Glc in KPC slice cultures, with total enrichments of 60% or greater by 48 hours (Fig. 3B, Extended Data Fig. 3A). In contrast, the labeling of pyrimidine nucleotides was much lower. UMP labeling remained under 20% in most tumors (Extended Data Fig. 3B), and CTP was essentially unlabeled through 96 hours of culture. Human PDAC tumor slices also exhibited this differential enrichment of purine and pyrimidine nucleotides (Fig. 3C). U-Glc can enrich both the ribose and nucleobase moieties of newly synthesized nucleotides, and ribose may also be labeled in the purine salvage pathway (Fig. 3D). The isotopologue distributions showed that AMP labeling from U-Glc was predominantly ribose labeling (M+5) in all tumors (Fig. 3E-F, Extended Data Fig. 3C), and ribose labeling was the predominant isotopologue detected in all nucleotides (Extended Data Fig. 3D-G). As shown in Fig. 2, glycine enrichment from U-Glc was low, thus we further quantified nucleotide synthesis using [^15^N] glutamine. Glutamine contributes nitrogen to purines and pyrimidines through both the α-amino and γ-amide groups (Extended Data Fig. 3H). Nearly 95% of the glutamine pool was enriched with ^15^N by either tracer within 48 hours, and α-Gln enriched 30-40% of the glutamate and aspartate pools (Extended Data Fig. 3I). Both α-Gln and γ-Gln labeled around 25% of the AMP pool in 48 hours (Extended Data Fig. 3J), much lower than the 60% enrichment from U-Glc. Consistent with the salvage of pyrimidine nucleosides, the UMP labeling from U-Glc and γ-Gln were comparable, and both were less than 30% at 48 hours. Together, these data suggest that *de novo* purine and pyrimidine synthesis do not contribute appreciably to maintaining nucleotide pools in PDAC slices. These results are consistent with findings in mice administered labeled nucleotides^47^, highlighting the general importance of salvage in maintaining nucleic acid homeostasis.

**Fig 3.**
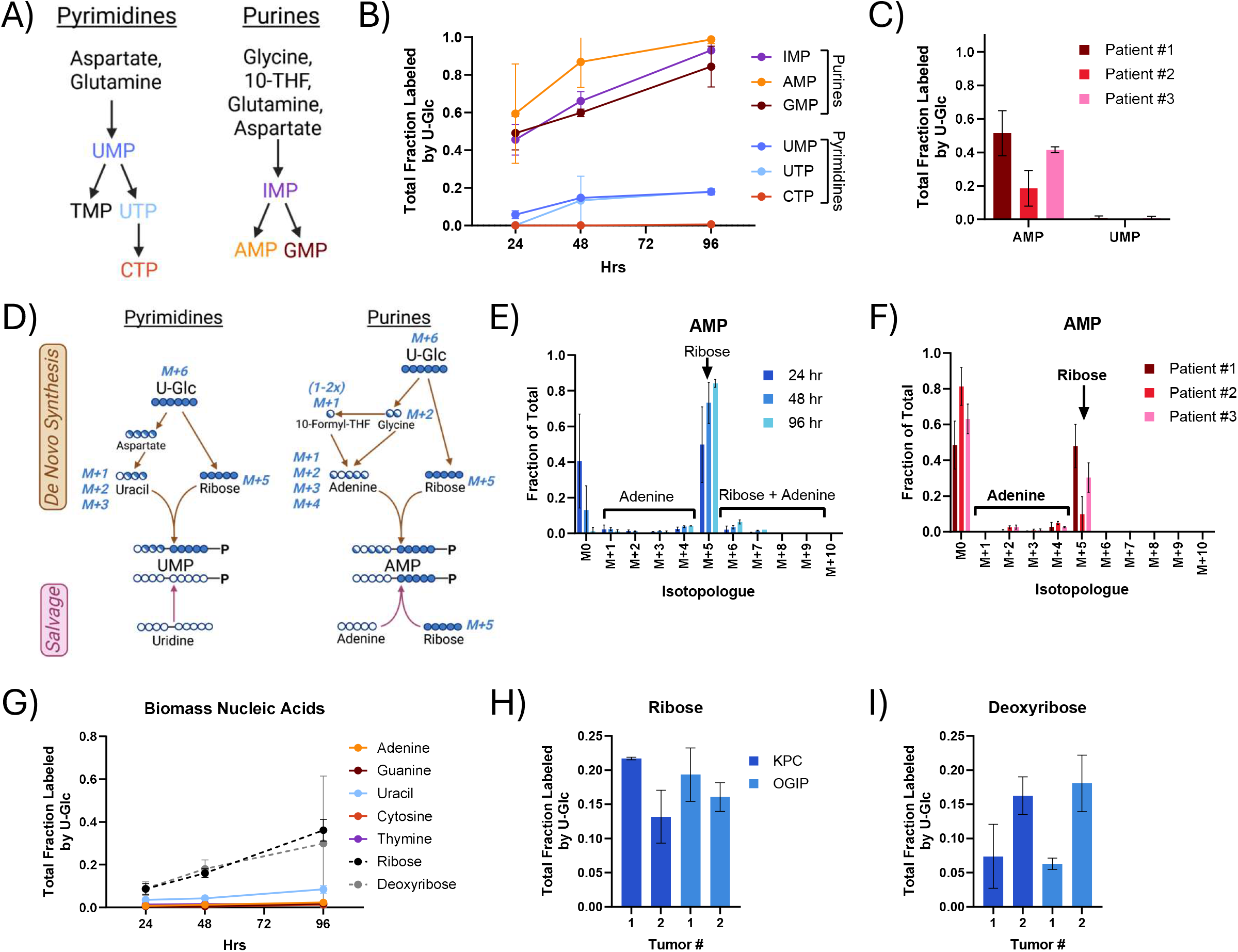
Purine and pyrimidine salvage supports nucleotide synthesis in PDAC tumor slices. (A) Diagram summarizing the *de novo* synthesis of pyrimidines and purines as nucleotide monophosphates. (B) Total labeling of the key nucleotides in *de novo* purine and pyrimidine synthesis identified in panel A via U-Glc tracing in tissue slices of KPC OGIP #2. TMP, TDP, or TTP were not quantifiable in PDAC slices from any KPC or KPC OGIP tumor. (C) Total fraction labeled from U-Glc of AMP and UMP in human PDAC slices at 48 hours. (D) Tracer map for possible labeling arising from U-Glc in pyrimidine and purine synthesis via *de novo* synthetic or salvage pathways. Isotopologue distribution of AMP from U-Glc in (E) tissue slices from KPC OGIP #2 and (F) human PDAC slice cultures at 48 hours. (G) Total labeling of nucleobases, ribose, and deoxyribose in the hydrolyzed biomass by U-Glc in tissue slices from OGIP #2. Total fraction labeled by U-Glc at 48 hours for (H) ribose and (I) deoxyribose in the hydrolyzed nucleic acids at 48 hours. All means represented three slices from a single tumor, and all error bars show standard deviation.

We also quantified enrichment of nucleobases, ribose, and deoxyribose within biomass via analysis of hydrolyzed macromolecular RNA and DNA. Consistent with the free nucleotide pools, the labeling of both purine and pyrimidine bases in RNA and DNA was low (Fig. 3G, Extended Data Fig. 3K). In contrast, the labeling of ribose and deoxyribose was higher and increased steadily through 96 hours, presumably via the incorporation of purine nucleotides enriched through the salvage pathway. Although the ribose moiety of the nucleotide was not 100%, these measurements also provide information on macromolecular nucleic acid synthesis within slices. At 48 hours, ribose and deoxyribose labeling was around 15-20% (Fig. 3H-I), similar to that of protein synthesis.

Additionally, nucleotides support glycosylation, and the glycosylation of proteins on the cell surface contributes to PDAC tumorigenesis and progression^48^. Nucleotide sugars, the active intermediates in glycosylation, were rapidly labeled by U-Glc in KPC and human PDAC slice cultures (Fig. 4A, Extended Data Fig. 4A). UDP-Hexose (UDP-Glucose and UDP-Galactose) was consistently nearly fully enriched by 24 hours. Nucleotide sugars can be labeled from U-Glc through multiple moieties (Fig. 4B), and UDP-Hexose was predominantly labeled on the activated hexose (M+6) (Fig. 4C-D) as anticipated from the low enrichment of the pyrimidine nucleotides. While less enriched overall, UDP-HexNAc (UDP-GlcNAc and UDP-GalNAc) was also consistently highly labeled in KPC and human PDAC slices (Extended Data Fig. 4B), primarily through the activated HexNAc (Extended Data Fig. 4C-F). CMP-SA labeling was distinct in KPC and human tumors, with human PDAC tumors having higher enrichment (Extended Data Fig. 4G). Notably, this labeling was likely via the sialic acid moiety in both tumor types (Extended Data Fig. 4H-J), although the isotopologue distribution of sialic acid may differ between species. Overall, this demonstrates that sugar interconversions are a major flux in PDAC tumor slices and that unlabeled nucleotides support nucleotide sugar synthesis.

**Fig 4.**
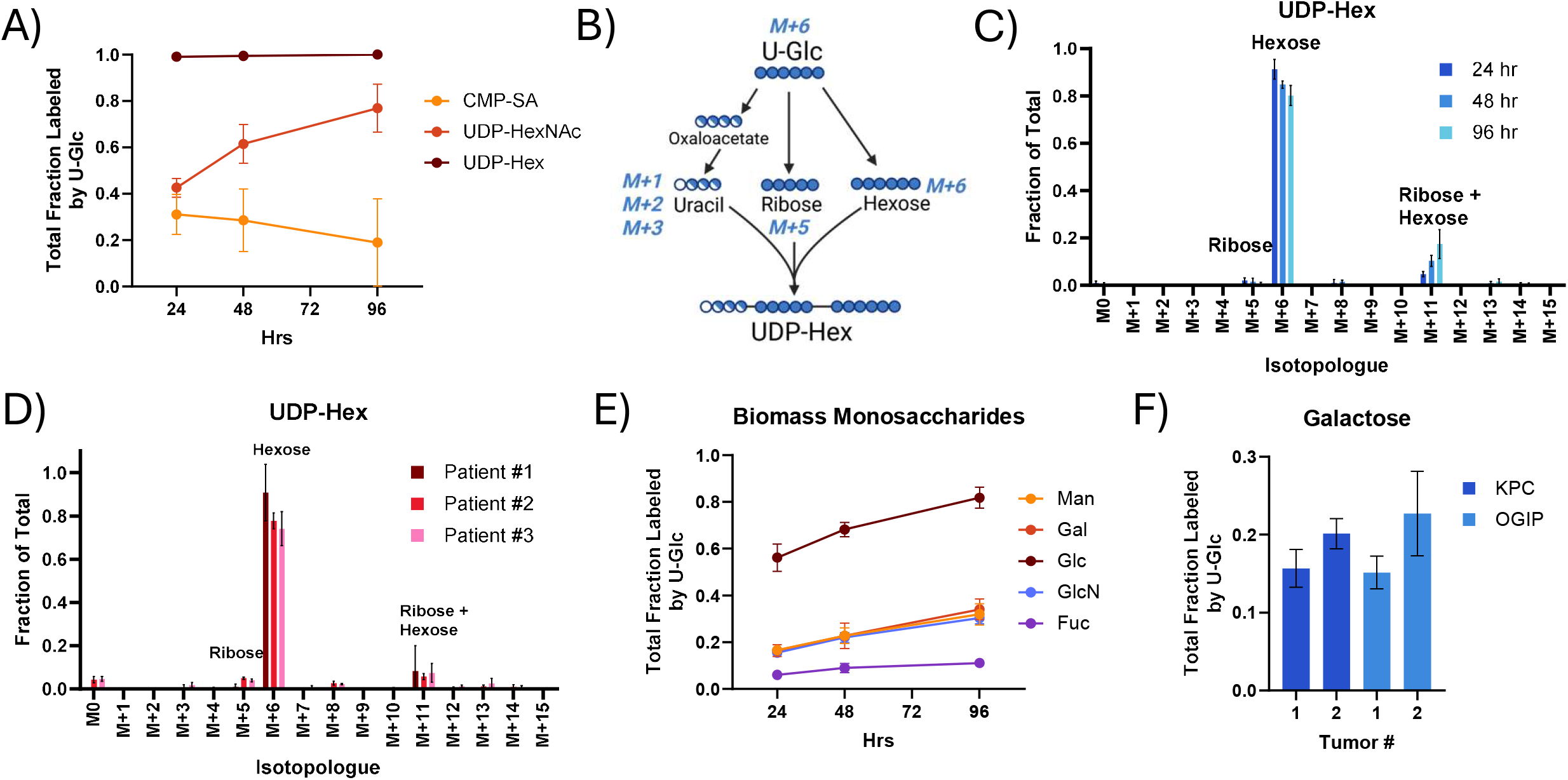
Nucleobase salvage and sugar interconversions maintain glycosylation flux in PDAC tumor slices. (A) Total fraction labeled by U-Glc for nucleotide sugars in tissue slices from KPC OGIP #2. UDP-glucuronic acid, GDP-mannose, and GDP-fucose were generally not present at quantifiable levels within PDAC slices. (B) Diagram demonstrating the possible labeling of UDP-Hexose (UDP-Hex; UDP-Glucose and UDP-Galactose) from U-Glc tracing. Isotopologue distribution of UDP-Hex from U-Glc in (C) tissue slices from KPC OGIP #2 and (D) human PDAC tumor slices at 48 hours. (E) Total labeling of sugars in the hydrolyzed biomass by U-Glc in tissue slices from OGIP #2. (F) Total labeling of galactose in the hydrolyzed biomass in KPC and KPC OGIP tumors at 48 hours . All means represented three slices from a single tumor. All error bars show standard deviation.

Acid hydrolysis of glycans from the biomass (i.e., proteoglycans and glycoproteins) revealed high synthesis rates in KPC tumor slices (Fig. 4E-F, Extended Data Fig. 4K). Labeling of glucose from biomass was very high, with greater than 60% enrichment at 48 hours. However, the supra-physiological concentration of glucose in cell culture media likely drives glycogen synthesis, as the abundance and labeling of glucose in the biomass were much more comparable to other monosaccharides when slices were cultured in 5 mM rather than 25 mM U-Glc (Extended Data Fig. 4L-M). Galactose, the other hexose constituent of the highly enriched UDP-Hexose pool, was around 15-25% labeled at 48 hours (Fig. 4F). Similarly, the glucosamine in the hydrolyzed biomass, representing both glucosamine and N-Acetyl-glucosamine, was around 20-30% labeled within 48 hours. This glycan labeling is higher than the 10-20% enrichment observed for proteins, RNA, and DNA synthesis within the same tumors at the same time point. Overall, the high labeling of monosaccharides in the biomass suggests that glycans are particularly highly synthesized macromolecules within the PDAC tumor slices and that the sugar interconversions maintaining glycosylation are a key flux of glucose within these tumors.

### Long-chain fatty acid salvage is critical in the PDAC tumor microenvironment

Citrate supplies lipogenic acetyl-CoA for *de novo* fatty acid synthesis. Despite consistently high citrate enrichment in KPC tumor slice cultures and human PDAC tumor slices (Extended Data Fig. 5A-B), we detected low labeling of saponified palmitate from U-Glc (Fig. 5A-B). Labeling of long-chain fatty acids (LCFAs) like palmitate, stearate, and oleate from U-Glc was consistently low across autochthonous and OGIP KPC tumors (Extended Data Fig. 5C). While other substrates can contribute to lipogenic acetyl-CoA pools, we detected very low labeling of LCFAs from ^13^C β-hydroxybutyrate, acetate, or glutamine in *ex vivo* slice cultures (Extended Data Fig. 5D-F). Taken together, these data establish that FASN flux is low in PDAC tumor slices.

**Fig 5.**
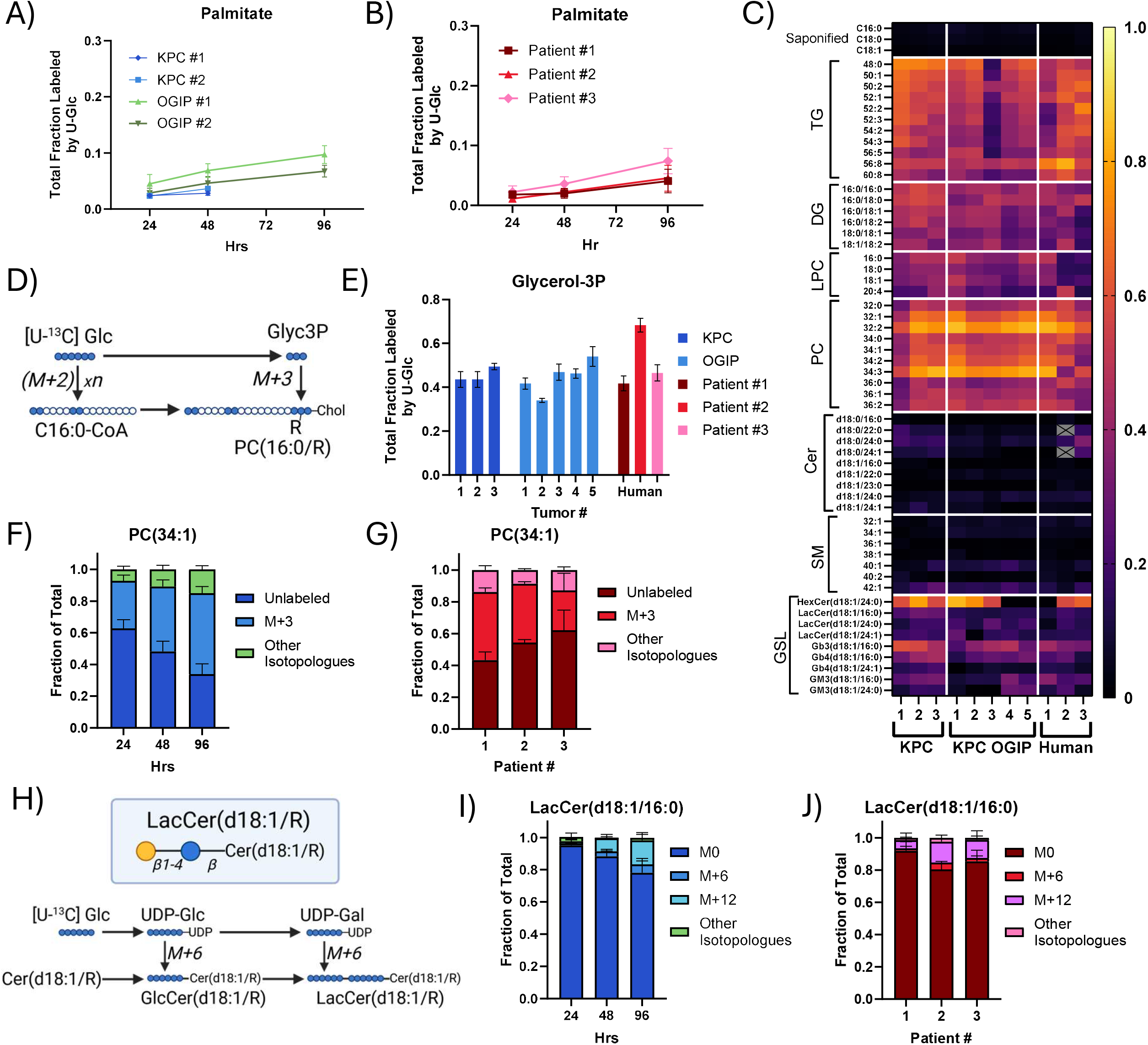
PDAC Tumor slices maintain high rates of lipid synthesis using salvaged long chain fatty acids. Total fraction labeled by U-Glc for palmitate in the saponified lipid fraction of (A) KPC tumor slices and (B) human PDAC tumor slices. (C) Heatmap of average total labeling of key and representative lipids from U-Glc at 48 hours in PDAC tumor slices. (TG, triacylglycerol; DG, diacylglycerol; LPC, lyso-phosphatidylcholine; PC, phosphatidylcholine; Cer, ceramide; SM, sphingomyelin; GSL, glycosphingolipid). Metabolites which were most abundant and most consistently quantifiable across tumors are shown. Metabolites which were not quantifiable within a particular tumor are shown in gray. (D) Diagram summarizing the synthesis and labeling of glycerolipids from glycerol 3-phosphate (Glyc3P) when tracing with U-Glc. (E) Total labeling of Glyc3P at 48 hours in PDAC tissue slices. Isotopologue distribution from U-Glc tracing into PC(34:1) in tissue slices from (F) KPC OGIP #2 and (G) human PDAC tumor slices at 48 hours. (H) Schematic demonstrating incorporation of labeled carbon from U-Glc into the sugar headgroups on GSLs like the lactosylceramides (LacCer). Isotopologue distribution of LacCer(d18:1/16:0) from U-Glc tissue slices from (I) KPC OGIP #2 and (J) human PDAC tumors at 48 hours. All means represented three slices from a single tumor. All error bars show standard deviation.

Complex lipids are major components of cell membranes required to support compartmentalization, cell signaling, and proliferation. We profiled diverse glycerolipid classes such as storage lipids (triacylglycerols; TGs, diacylglycerols; DGs) and key membrane lipids (lysophosphatidylcholines; LPCs, phosphatidylcholines; PCs) after 48 hours of tracing with U-Glc. Unexpectedly, the overall enrichment of most glycerolipids was much higher than fatty acid synthesis rates in both KPC and human PDAC tumor slices (Fig. 5C), with enrichments of 40-60%. These glycerolipids are comprised of fatty acid acyl-chains on a glycerol 3-phosphate (Glyc3P) headgroup backbone. While U-Glc derived acetyl-CoA incorporates labeled carbon from U-Glc in units of two, Glyc3P directly incorporates three labeled carbons (M+3) into lipid headgroups (Fig. 5D). Unlike fatty acids, U-Glc efficiently labels Glyc3P in slice cultures (Fig. 5E), and the isotopologue distribution for both glycerophospholipids and TGs revealed high M+3 enrichment (Fig. 5F-G, Extended Data Fig. 5G-J), indicative of Glyc3P backbone labeling. Lipids containing very long-chain fatty acid (VLCFA) acyl chains, such as LPC(24:0), also exhibited appreciable labeling derived from the elongation of LCFAs via ELOVL elongases^49^, as evidenced by the incorporating 2-carbon labeled acetyl-CoA units from U-Glc to form M+5, M+7, and M+9 isotopologues (Extended Data Fig. 4K-L). Intriguingly, glycerolipids and the Glyc3P pool often show similar labeling between 24 and 48 hours (Extended Data Fig. 5M), indicating these metabolite pools rapidly achieve isotopic steady state. In other words, nearly the entire pool of glycerolipids within these slices was (re)synthesized over the 48-hour experiment. Lands cycle-associated reactions are a major contributor to this turnover^50^ but the high rates of labeled Glyc3P incorporation confirm other pathways are involved as well. In contrast, we observed around 10-20% protein and nucleic acid turnover during the same period. These results indicate that glycerolipid turnover is high within tumors and suggest that lipid remodeling is an important aspect of tumor cell growth and survival within the PDAC TME.

While sphingolipids like ceramides and sphingomyelins (SMs) were generally less than 20% labeled by U-Glc at 48 hours (Fig. 5C), many glycan-bearing glycosphingolipids (GSLs) were 30-50% labeled. Serine, which provides the headgroup backbone for sphingolipids, was only around 10-20% enriched from U-Glc (Extended Data Fig. 6A). Thus, low enrichment of both serine and fatty acid pools by U-Glc limits the labeling of non-glycosylated sphingolipids like SMs, particularly in human PDAC slices (Extended Data Fig. 6B-C). In contrast, the isotopologue distribution of GSLs was indicative of sugar headgroup labeling, such as M+6 and M+12 labeling on lactosylceramide (LacCer) (Fig. 5H-J, Extended Data Fig. 6D) and M+18 labeling on the globoside Gb3 (Extended Data Fig. 6E-F). In tissue slices from Patient #1, [U-^13^C] Serine (U-Ser) labeled dihydroceramides, a key intermediate in *de novo* sphingolipid synthesis, and many GSLs by 10-20% in 24 hours (Extended Data Fig. 6G-H), providing evidence that *de novo* sphingolipid synthesis is active within human PDAC tumors. Notably, many GSLs were more enriched by U-Glc on the sugar headgroup than U-Ser on the lipid moiety, especially the globoside Gb3 and the ganglioside GM3(d18:1/16:0), which indicates ceramide salvage. Additionally, ceramides and SMs with VLCFA acyl chains were more highly enriched than LCFA-containing species (Fig. 5C), with isotopologue distributions consistent with labeling predominantly via chain elongation (Extended Data Fig. 6I-J). Together, these data highlight the importance of headgroup turnover and ELOVL activity rather than LCFA synthesis in sustaining GSL pools in both KPC and human PDAC slices.

### Systemic lipid metabolism supplies local lipid availability for LCFA salvage

As PDAC tumors within the native TME maintained high rates of lipid synthesis without high rates of *de novo* fatty acid synthesis, we next investigated the role of the TME in lipid homeostasis by comparing 2D KPC-derived FC1245 cells to *ex vivo* slices of orthotopic tumors generated with the same cell line in syngeneic mice (Fig. 6A). While PDAC cells were highly lipogenic in 2D culture, palmitate synthesis was dramatically lower in slice cultures from orthotopic tumors (Fig. 6B) and was equivalent to rates found in KPC, KPC OGIP, and human PDAC tumor slices (Fig. 5A-B). FASN flux in the orthotopic tumors was particularly reduced despite the high contribution of U-Glc to the lipogenic acetyl-CoA pool through citrate (Extended Data Fig. 7A-B). This phenotype was not specific to the FC1245 cell line or to murine PDAC cells, as Suit2 cells experienced a similar decrease in *de novo* lipogenesis when cultured as tissue slices of xenografted tumors (Extended Data Fig. 7C). Together, these data demonstrate that while cells in both models had high rates of lipid turnover, FASN flux was only active in 2D culture. Thus, the *in situ* TME, rather than adaptation to 2D culture, primarily determines the reliance on FASN-mediated *de novo* lipogenesis.

**Fig 6.**
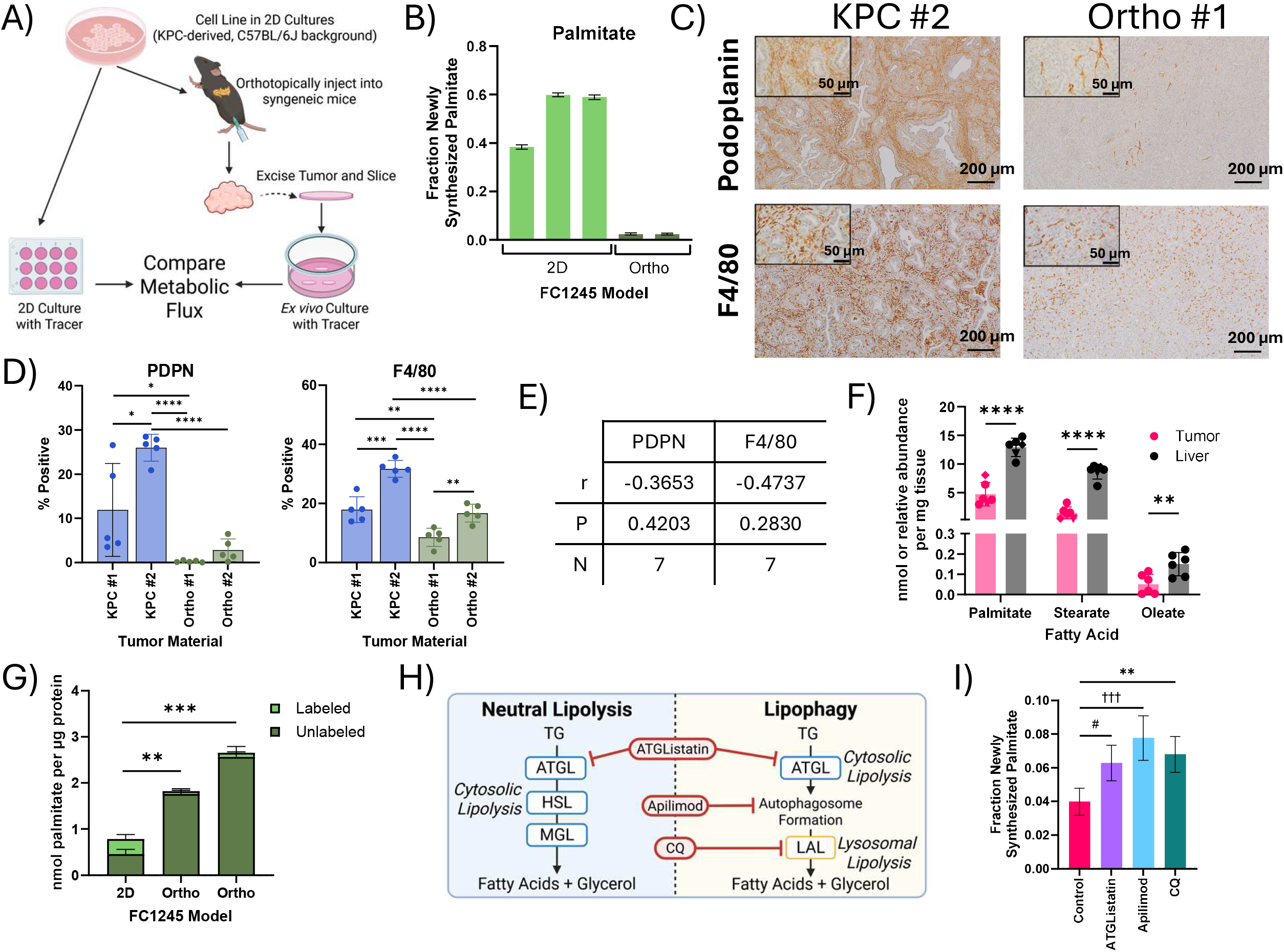
Local lipid availability within the PDAC tumor microenvironment maintains lipid synthesis through salvage. (A) Schematic describing the design of metabolic flux studies comparing baseline lipid synthesis of PDAC cells in 2D culture and orthotopic tumors. (B) Palmitate synthesis in FC1245 cells in 2D culture and in tissue slices from orthotopic tumors (Orthos), calculated using isotopomer spectral analysis (ISA) of U-Glc labeling of palmitate in the saponified fatty acids. Error bars represent 95% confidence intervals. (C) Immunohistochemistry staining for podoplanin or F4/80 in primary tumor material from a KPC tumor and FC1242 Ortho. Scale bars represent 200 μm, inset scale bars are 50 μm. (D) Quantitation of the percent of cells positive for podoplanin (PDPN) and F4/80 in primary tumor material from KPC tumors and FC1242 Orthos. Data points represent independent fields of view (n=5). (E) Pearson correlation between the total labeling of palmitate at 48 hours and the percentage of cells positive for PDPN or F4/80 in primary tumor material. (F) Newly synthesized long chain fatty acids in KPC OGIP tumors and matched liver tissue (n=6). Quantitation was performed following 48 hours of *in vivo* D_2_O tracing and analyzed from the saponified lipid fraction. (G) Palmitate abundance in the saponified lipids from FC1245 Ortho tumor slices and 2D cultures after 48 hours of culture with U-Glc. (H) Diagram summarizing the lipolysis of triacylglycerols (TGs) to release fatty acids (ATGL, adipose triglyceride lipase; HSL, hormone sensitive lipase; MGL, monoglyceride lipase; LAL, lysosomal acid lipase (LAL) and inhibitors of these steps (CQ, chloroquine). (I) Palmitate synthesis with 10 μM ATGListatin, 100 nM apilimod, or 25 μM CQ for 48 hours by ISA modeling of the saponified lipid fraction in tissue slices from KPC OGIP #5. Error bars represent 95% confidence intervals. Statistical significance was determined by overlapping error bars and denoted as # for ATGListatin, † for Apilimod, * for CQ. Unless otherwise specified, means represented three slices from a single tumor or three wells of 2D cell culture, and error bars show standard deviation. Where relevant, statistical significance was determined by two-tailed *t*-test (*, *P* < 0.05; **, *P* < 0.01; ***, *P* < 0.001; ****, *P* < 0.0001).

The comparability of *de novo* lipogenesis in KPC and 2D-derived orthotopic tumors was striking (Extended Data Fig. 7D), as orthotopic tumors from 2D cell lines do not fully recapitulate the stromal diversity observed in autochthonous and OGIP KPC tumors^44^. 2D-derived FC1242 orthotopic tumors were more homogenous and exhibited lower infiltration of fibroblasts and macrophages than autochthonous KPC tumors (Fig. 6C-D, Extended Data Fig. 7E). We observed no significant correlation between the proportion of fibroblasts (podoplanin, PDPN) or macrophages (F4/80) and palmitate synthesis at 24 or 48 hours (Fig. 6E, Extended Data Fig. 7F). Since fatty acid synthesis within these different PDAC tumor slices was not significantly impacted by tumor cellularity, these data suggest that fatty acid salvage is supported by both cellular and non-cellular elements of the PDAC TME.

Lipid availability is notably different between 2D cell culture conditions and *in vivo*^9,51,52^. To compare our results in slice culture to *in vivo* tumor growth conditions, we next administered D_2_O to mice bearing KPC OGIP tumors for 48 hours to quantify the abundance of newly synthesized palmitate in liver and tumor tissue. While newly synthesized palmitate was present in all tumors, the abundance was significantly lower than that observed in the liver (Fig. 6F), suggesting that hepatic lipogenesis and lipoprotein transport supplies tumors with fatty acids *in vivo*. Indeed, orthotopic tumors had around two-fold more palmitate per μg protein compared to 2D culture (Fig. 6G), highlighting local availability from stroma and tumor interstitial fluid. Many cells, including PDAC cells, will readily scavenge extracellular lipids from serum^14^, and culturing 2D cells in 20% FBS has been reported to more accurately recapitulated the lipid synthesis observed in *ex vivo* tissue slice cultures from murine lung cancer models^52^. However, increasing the medium FBS concentration to 20% only modestly decreased the contributions of palmitate synthesis in 2D cultures of PDAC cells (Extended Data Fig. 7G). Moreover, fatty acid synthesis in KPC slices was not sensitive to serum concentration (Extended Data Fig. 7H). Thus, these data suggest that exposure to systemic lipid metabolism rather than tumor cellularity determines the role of LCFA salvage in PDAC cells.

Fatty acids are distributed systemically as TGs via lipoproteins and are released from these transported lipoproteins and locally-stored TGs within lipid droplets through lipolysis^53,54^. This process contributes to the proliferation and metastasis of PDAC cells^55,56^. An advantage of slice culture is the ability to test multiple pharmaceutical agents within the same tumor, so we next employed MFA to assess the impact of different small molecule inhibitors of lipid salvage pathways on membrane lipid homeostasis. Free fatty acids are mobilized from TGs via Adipose Triglyceride Lipase (ATGL)-mediated neutral lipolysis or via lysosomal acid lipase (LAL) during lipophagy^53,56^. The lipid kinase PIKfyve is an established coordinator of endolysosomal trafficking and autophagic flux^57^ which was recently shown to impact lipid metabolism in PDAC tumors^55^. When we inhibited neutral lipolysis using ATGListatin, or lysosomal degradation using apilimod or chloroquine (CQ) (Fig. 6H, Extended Data Fig. 7I), total palmitate synthesis increased by nearly two-fold in KPC OGIP tissue slices (Fig. 6I). While reinforcing our findings that salvage is the predominant source of fatty acids in the PDAC TMC, there was no significant impact on saponified fatty acid or lipid abundances in slice cultures with any inhibitor (Extended Data Fig. 7J). This highlights the propensity for metabolic compensation within the PDAC TME and demonstrates the utility of MFA in slice culture for assessing molecular mechanisms supporting lipid homeostasis.

Despite having similar impacts on FASN flux, these inhibitors had notably different impacts on downstream membrane lipids. While none of the inhibitors significantly impacted the abundance or overall labeling of TGs within the slices after 48 hours, ATGListatin and apilimod significantly increased acyl chain labeling (Extended Data Fig. 7K-M). In contrast, apilimod and CQ, but not ATGListatin, perturbed the overall labeling of DGs and glycerophospholipids (Extended Data Fig. 7N). While LPC labeling was particularly impacted, neither apilimod nor CQ affected the abundance of LPCs (Extended Data Fig. 7O), further highlighting the compensatory capabilities of PDAC slices and the importance of applying MFA to assess metabolic responses.

### PIKfyve supports ganglioside synthesis in PDAC cells

PDAC tumor slices actively re-synthesize complex GSLs like gangliosides and globosides using salvaged fatty acids and ceramides (Fig. 5). Unlike fatty acid synthesis, GSL diversity and abundance was reflective of tumor cellularity within multiple models of PDAC (Extended Data Fig. 8A-C). PDAC cells are known to express gangliosides, especially GM2^58–60^, and gangliosides were much more abundant than globosides in 2D cultures of “pure” PDAC cells (Fig. 7A). FC1245 cells expressed diverse gangliosides but had few quantifiable globosides. FC1245 orthotopic tumors, in which we observed only modest stromal infiltration (Fig. 6D-E), expressed similarly diverse gangliosides with greater globoside abundance. Finally, autochthonous KPC tumor slices exhibited diverse gangliosides and globosides, with the non-sialylated globosides being significantly more abundant in these stroma-enriched tumors. Single cell RNA-Seq data from KPC tumors on key enzymes in this pathway highlight distinctions in expression between ductal cells and fibroblasts (Fig. 7C). Notably, many enzymes involved in GSL synthesis are expressed by multiple cell types within the TME, including UDP-glucose ceramide glucosyltransferase (*Ugcg*), the gene encoding the common first step in both ganglioside and globoside synthesis. On the other hand, GM3 synthase (*St3gal5*) and Gb3 synthase (*A4galt*) are much more narrowly expressed. GM3 synthase is primarily expressed in ductal tumor cells, endothelial cells, and mesothelial cells. As endothelial and mesothelial cells only comprise a small proportion of the tumor (Extended Data Fig. 8D), this suggests that GM3 within these slice cultures is primarily synthesized by tumor cells. Conversely, the expression of globoside synthesis enzymes like Gb3 synthase is limited to endothelial cells and fibroblasts, indicating that fibroblasts may be the predominant source of globosides within the PDAC TME. PDAC tumors have been shown to contain diverse GSLs ^61^, including both gangliosides and globosides. These data suggest that GSL biosynthesis is intrinsic to diverse cells within PDAC tumors and that these processes are coordinated across the TME.

**Fig 7.**
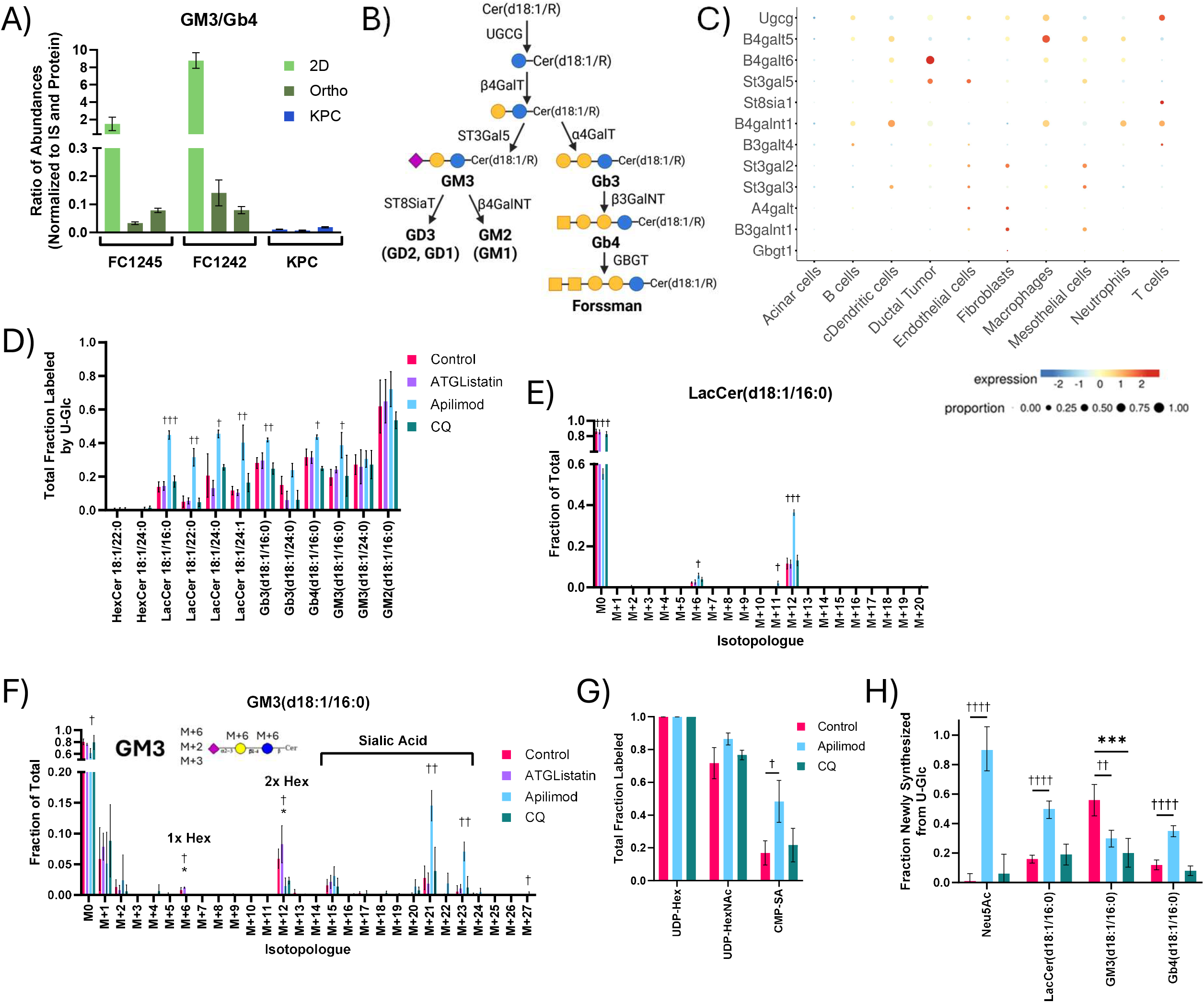
PIKfyve supports glycosphingolipid synthesis in tumor and stromal cells. (A) Ratio of the abundance of GM3(d18:1/16:0) to Gb4(d18:1/16:0) at 24 hours across multiple models of PDAC. These were selected as representative ganglioside and globosides containing the same lipid moiety which were quantifiable across all models. (B) Schematic depicting the key enzymes involved in the synthesis of gangliosides and globosides. (C) Proportion and relative expression of key glycosphingolipid synthesis genes by single cell RNASeq analysis of KPC tumors (GSE125588). (D) Total labeling of glycosphingolipids from U-Glc after 48 hours of tracing with 10 μM ATGListatin, 100 nM apilimod, or 25 μM CQ in KPC OGIP #5. (E) LacCer(d18:1/16:0) and (F) GM3(d18:1/16:0) isotopologue distributions with inhibitor treatments. (G) Total labeling of nucleotide sugars from U-Glc at 48 hours. (H) Synthesis fluxes for n-acetyl-neuraminic acid and key glycosphingolipid species as calculated by Lipid Metabolic Flux Analysis (Lipid MFA) modeling of isotopologue distributions following 48 hours of U-Glc tracing. Error bars represent 95% confidence intervals, and statistic significance is determined by overlapping error bars. All means represented three slices from a single tumor or three wells of 2D cell culture. Unless otherwise specified, error bars show standard deviation, and statistical significance was determined by two-tailed *t*-test. Significance denoted as # for ATGListatin, † for Apilimod, * for CQ. (*, *P* < 0.05; **, *P* < 0.01; ***, *P* < 0.001; ****, *P* < 0.0001).

Due to the observed contributions of cellular diversity to GSLs, studying these pathways within an intact TME is critical to understanding how the pathway may be remodeled in the context of therapy. Intriguingly, apilimod perturbed GSL enrichment more than any other treatment. Apilimod dramatically increased overall labeling of most GSLs in slice culture (Fig. 7D) while abundances were maintained (Extended Data Fig. 8E), indicating cells compensate for defects in salvage with greater synthesis to maintain GSL pools. Interestingly, while apilimod also significantly increased acyl chain labeling on ceramides and sphingomyelins to maintain abundances (Extended Data Fig. 8F-H), increased labeling of the GSLs was almost exclusively on the sugar headgroups rather than the lipid moieties (Fig. 7E-F). The labeling of sialic acid-containing ganglioside GM3 was particularly increased by apilimod treatment in multiple tumors, particularly via the proportion of isotopologues representing sugar headgroups with newly synthesized sialic acid (i.e., M+21 and M+23) (Fig. 7F, Extended Data Fig. 8I). Apilimod also significantly increased the labeling of the sialic acid moiety on CMP-sialic acid (Fig. 7G, Extended Data Fig. 8J), suggesting that PIKfyve may play a role in sialic acid metabolism. To deconvolute the impacts on sugar and lipid metabolism in GSLs, we employed Lipid MFA modeling using a network of the most impacted GSLs: LacCer(d18:1/16:0), GM3(d18:1/16:0), and Gb4(d18:1/16:0) (Extended Data Fig. 8K). Apilimod dramatically increased the contribution of newly synthesized sialic acid to GM3 while the overall synthesis of GM3 was significantly decreased (Fig. 7H). This provides evidence that targeting PIKfyve impacts both glycan and GSL homeostasis in PDAC tumor cells. While CQ treatment similarly decreased GM3 synthesis, sialic acid synthesis was not impacted, suggesting that the role of PIKfyve in sialic acid metabolism may be independent of the lysosome. LacCer and Gb4 synthesis were also significantly impacted by apilimod, revealing that PIKfyve may play a role in GSL metabolism in fibroblasts as well. However, the contrasting impacts of apilimod on ganglioside and globoside metabolism provide evidence that PIKfyve may support GSL metabolism through distinct mechanisms in PDAC tumor cells and fibroblasts. Given the importance of GSLs in establishing cell identity, these data highlight the role of PIKfyve-mediated phosphoinositol signaling and lipid salvage fluxes in coordinating the membrane lipid and glycan homeostasis of PDAC within the native TME.

## Discussion

In this paper, we quantify the synthesis of proteins, complex lipids, nucleic acids, and glycans in various PDAC models, including 2D cell cultures and *ex vivo* slice cultures that retain an intact TME. Through the analysis of multiple murine and human tumors, we determined that nucleobases and LCFAs were predominantly salvaged when evaluated within the PDAC TME. This reliance on LCFA salvage was particularly striking, as PDAC tumor slices exhibited very high rates of glycerolipid turnover relative to other macromolecules. While appreciable patient-to-patient variability has been demonstrated in PDAC^64,65^ every tumor we investigated demonstrated this reliance on LCFA salvage, including multiple human tumors and xenografts generated from Suit2 cells, an established lipogenic cell line^64^. Because appreciable FASN flux occurs in tissue slices from mice bearing non-small cell lung carcinomas^52^, our results suggest LCFA salvage may be intrinsic to PDAC. Indeed, PDAC cells cultured in 2D display a high rate of FASN flux that is diminished in the context of slice culture. Intriguingly, PDAC tumors are generally considered to have elevated expression of and reliance on FASN compared to healthy tissues^65–68^, and FASN-mediated lipogenesis plays an established role in PDAC cell culture^62,68,69^. The role of FASN in PDAC remains somewhat unclear^70^, and FASN expression in PDAC may be serving other purposes beyond bulk lipid synthesis, including palmitoylation to stabilize mutant p53^71^ or regulation of DNA repair activity through NF-κB signaling^72^. However, more direct interrogation of these substrates is needed to explore potential non-metabolic functions of FASN.

Rather than rely on local LCFA synthesis, these data suggest that PDAC tumors employ multiple lipid salvage pathways for provision of membrane lipid precursors within the PDAC TME, including lipoprotein transport, neutral lipolysis, and lipophagy^56,73,74^. These pathways ultimately fuel the synthesis of tumor-intrinsic gangliosides like GM3 which have diverse functions in PDAC cells such as maintaining Kras localization^60^, modulating immune interactions^75,76^, and promoting metastasis^77^. Notably, the PIKfyve inhibitor apilimod perturbs GSL synthesis through modulating both sialic acid homeostasis and lipid metabolism, suggesting that PIKfyve supports the role of gangliosides in PDAC tumor progression^58,62^. PIKfyve is thought to have both lysosomal and non-lysosomal functions in retrograde transport between the endosome and the trans-Golgi network^57,78,79^. Our direct comparison of apilimod and CQ, a known modulator of lysosomal lipid salvage, suggests that non-lysosomal functions of PIKfyve may be particularly relevant for sialic acid metabolism. Sialic acid homeostasis is also critical for the synthesis of other glycans, including sialyl Lewis A epitope (CA19-9), which is upregulated in PDAC and promotes tumorigenesis and disease progression^80^. However, our results also highlight key redundancies that enable PDAC cells within the TME to adapt and sustain provision of lipids when these salvage pathways are perturbed, particularly increased reliance on *de novo* synthesis of fatty acids and sialic acid. These compensatory pathways may present potential co-targets and/or mechanisms of resistance.

Although the limited amount of human PDAC tumor material available for research can be restrictive for other applications like pharmacotyping^32–38^, slice cultures and MFA provide a model in which these metabolic signatures can be identified using relatively little tumor material. Indeed, the ability to avoid tumor-to-tumor variability by directly employing multiple stable isotope tracers or small molecule inhibitors on the same tumor is a key advantage of this MFA approach. Labeling changes and fluxes are likely more informative than pool sizes, which changed minimally in response to treatments due to pathway redundancies. However, flux data are currently limited to the bulk metabolism of the tumor slices, which encompass the metabolic fluxes of both tumor cells and stromal cells. Although these fluxes may predominantly represent the contributions of a particular cell type, as we observed in GSL synthesis, we cannot formally elucidate the fluxes within tumor or stromal cells specifically using bulk tissue metabolomics alone. Advancements in spatial-oriented analytical technologies, such as spatial mass spectrometry, may soon enable resolution of metabolic fluxes on a cellular level^81^, thus enabling the elucidation of cell-type specific fluxes within PDAC tumors. Finally, the ability to employ pharmacological inhibitors or chemotherapeutic agents in a patient’s own tumor *ex vivo* may provide mechanistic information on tumor cell growth and survival pathways that aids in the identification of novel metabolic targets to enhance therapeutic efficacy or combat resistance as well as enable testing for personalized responses to specific therapeutic agents.

## Acknowledgements

We thank all members of the Metallo lab for support, especially Karl Wessendorf-Rodríguez for providing guidance on the flux modeling and Emily Fennell for providing feedback on this manuscript. We also thank members of the Engle lab for their insights and support, especially Vasiliki Pantazopoulou.

We acknowledge support from NIH grant R01CA234245 (to C.M.M.), NIH Grant UO1CA274295 (to A.M.L.), the Mark Foundation for Cancer Research (to C.M.M.), the Salk NCI Cancer Center CCSG P30 CA013195, the National Cancer Institute P01 CA265762 (to D.E.E. and A.M.L), the Lustgarten Foundation 122215393, the Paul M. Angell Family Foundation SA-SP24 (to D.D.E.), the American Cancer Society 1252509, and the Conrad Prebys Foundation Excellence in Scientific Research Leadership Program. The NIH grants T32CA009370 (to C.S.K.)

The authors wish to acknowledge the support of the Moores Cancer Center Biorepository and Tissue Technology Shared Resource, supported by the National Cancer Institute of the National Institutes of Health under award number P30CA23100. The content is solely the responsibility of the authors and does not necessarily represent the official views of the National Institutes of Health.

Figures created using GraphPad Prism10 and Biorender (Biorender.com).

## Methods

### Animal Studies

All experiments were conducted in accordance with the Institutional Animal Care and Use Committee (IACUC) of the Salk Institute for Biological Studies.

For orthotopically grafted intact pieces (OGIP) generation, tumor-bearing KPC mice were sacrificed and the tumor dissected. Each tumor was cut into 2-3 mm^3^ pieces, washed, and cryopreserved in liquid nitrogen for future use. To generate KPC OGIP tumors, OGIPs were thawed on ice. Mice were anesthetized via isoflurane inhalation, hair at the surgical site was removed by plucking, and the outer skin disinfected. After verifying adequate sedation via toe pinch, the pancreas was exteriorized through a small left flank incision and an OGIP was wrapped into the tail of the pancreas and secured using tissue glue. The peritoneum was then sutured closed, and the skin was shut with wound clips.

For orthotopic tumors generated from 2D cell culture, cells were dissociated with 0.25% trypsin-EDTA, then suspended in 50% DMEM + 50% BME. For KPC-derived orthotopic tumors, 1,000 FC1245 cells or 10,000 FC1242 cells/mouse were injected into the tail of the pancreas of female or male C57BL/6 mice, respectively. For the Suit2 xenografts, 30,000 Suit2 cells/mouse were orthotopically injected 8 week old male NSG mice. The peritoneum was then sutured closed, and the skin was shut with wound clips. After OGIP and orthotopic surgery, all mice were given 1mg/kg of sustained released buprenorphine as an analgesic.

Tumor growth was monitored by weekly ultrasound. Mice were euthanized prior to humane endpoint or earlier based on study-specific criteria. Mice bearing Suit2 xenografts were enrolled onto isocaloric diets containing either 10% (LFD) or 60% (HFD) fat when tumors reached 4 mm in diameter and maintained on these diets for 3 weeks prior to euthanasia. Mice were euthanized by cardiac puncture and cervical dislocation. All tissues were collected in the fed state, unless otherwise noted.

### D_2_O Tracing for quantification of fatty acid synthesis

Male C57BL/6 mice bearing KPC OGIP tumors 7-9 mm in diameter were administered ^2^H2O (D_2_O) in 0.9% NaCl via intraperitoneal injection at a dose of 0.027 mL/g body weight. 8% D_2_O drinking water was provided ad libitum. Mice were fasted for 6 hours prior to euthanasia, and 8% D_2_O drinking water was provided ad libitum during the fasting period. 48 hours after injection, mice were euthanized by cardiac puncture. Plasma was collected as previously described^82–84^. Tissues were snap-frozen in LN_2_ using pre-cooled Wollenberger clamps and stored at -80 ^o^C until analysis.

D_2_O enrichment of fatty acids in liver and tumor were quantified as previously described^82–84^. Briefly, 10-20 mg of liver or tumor tissue was homogenized in 1 mL of cold 1:1:2 MeOH:H_2_O:CHCl_3_ with 1 ug/sample palmitate-d_31_ internal standard. The CHCl_3_ phase was collected, dried under N_2_, and stored at –80 ^o^C for fatty acid analysis via gas chromatography–mass spectrometry (GC–MS) as described below. Fatty acid synthesis was normalized to plasma D_2_O enrichment via an external standard curve using the deuterium–acetone exchange protocol ^83,84^. The fraction of newly synthesized fatty acids was calculated according to previously published methodology ^85^.

### Tissue Slicing

Upon sacrifice, murine tumors were removed, weighed, and length and width measured. Pieces of tumor were quickly removed for histological staining; the remaining tumor was placed in ice cold Krebs-Henseleit buffer. Smaller tumors were superglued to a stage for vibratome sectioning, and larger tumors were cut into pieces prior to gluing. 200μm sections were cut on a Leica VT1200S with Vibracheck. Sections were kept continuously in chilled Krebs-Henseleit buffer prior to plating.

For human PDAC material, tissues were obtained from patients undergoing surgical resection or tissue biopsy at the University of California San Diego Health. All tissue donations and experiments were reviewed and approved by the Institutional Review Board of the University of California San Diego Health. Written informed consent was obtained prior to the acquisition of tissue from all patients. Tissue slice was performed as previously described ^36,86^. Briefly, tissue material was divided into sections and embedded into 4% low-gelling temperature agarose in PBS. 200μm sections were cut on a Leica VT1000P.

### Tumor Slice Culture and Stable Isotope Tracing

PDAC tissue slices are cultured *ex vivo* as previously described ^40,87^. Briefly, three tissue slices are transferred into pre-equilibrated Recovery Media (RPMI 1640 + 2% fetal bovine serum (FBS) + 1x penicillin/streptomycin (P/S)) in a 6 well plate. Transwell inserts (24 mm, 8.0 um polycarbonate) were used for agarose embedded human tissue, and low attachment suspension culture plates were used for non-embedded murine slices. Care was taken to ensure no overlap or contact between slices. To address tumor heterogeneity and avoid biasing based on tumor region, slices were selected from the same or matching tumor regions, and tissue slices were plated sequentially across the compared conditions to more evenly distribute intra-region variability. Tissue slices were cultured in a humidified incubator at 37 ^o^C and 5% CO_2_.

Tissue slices were allowed to recover for at least 16 hours overnight prior to tracing studies. Wells were washed once with sterile dPBS, then replace with tracing medium at 3 mL/ well in low attachment plate or 5 mL/well with transwell inserts. Tracing medium for murine tumor slices was formulated using Dulbecco’s Modified Eagle’s Medium (DMEM) without Glucose or Glutamine supplemented with 2% FBS and 1% P/S. For ^13^C tracing studies, 5 mM or 25 mM [U-^13^C_6_] D-Glucose or 4 mM [U-^13^C_5_] L-Glutamine were added to the tracing medium, as required for the study design. ^12^C glucose or glutamine were then added to target 25 mM glucose or 4 mM Glutamine, as needed. For tracing with U-βHB or U-Acetate, 1 mM [U-^13^C_4_] beta-hydroxybutyrate or 1 mM [1,2-^13^C_2_] Acetate were supplemented into DMEM with 2% FBS and 1% P/S. For ^15^N tracing, DMEM without Glucose or Glutamine was supplemented with 5 mM ^12^C glucose, 2% FBS, 1% P/S, and either 4 mM [α-^15^N] Glutamine or 4 mM [amide-^15^N] Glutamine. Tracing medium for human tissue slice studies, including Suit2 xenografts, was formulated using RPMI 1640 without Glucose or RPMI 1640 without glucose or amino acids. [U-^13^C_6_] Glucose or [U-^13^C_3_] Serine were added at concentrations consistent with RPMI 1640 medium, and remaining glucose and amino acids, 2% FBS, and 1% P/S were added. For tracing studies lasting longer than 24 hours, tracing medium was exchanged daily. At desired time points, tracing medium was removed, and slices in wells were washed twice with 0.9% saline. Tissue slices were placed in Eppendorf tubes individually, flash frozen in LN2, and stored at -80 ^o^C until extraction.

For inhibitor studies, inhibitors were formulated in the appropriate vehicle, then added directly to the tracer medium at the target concentration for all exchanges. Inhibitor stock solution concentrations and vehicles were as follows: Chloroquine Diphosphate (15 mM in dPBS), ATGListatin (10 mM in DMSO), Apilimod (100 μM in DMSO). Target concentrations are as described in text and figure legends.

### Cell culture studies

FC1242, FC1245, and Suit2 cell lines were obtained from the Engle Lab. Cell lines were periodically tested for and found to be free of mycoplasma contamination. All 2D cultures were cultured in DMEM with 10% FBS and 1x P/S for maintenance and expansion unless otherwise noted.

To conduct stable isotope tracing studies, cells were plated at 100,000 cells/well in a 12 well tissue culture plate or 150,000 cells/well in a 6 well tissue culture plate. For consistency between tissue slice cultures and 2D culture, cells were plated into Recovery Medium. After 16-24 hours, wells were washed once with sterile dPBS, and media was replaced with 1 mL/well tracing medium. Tracing medium was prepared as previously described for tissue slice cultures and was exchanged daily.

Metabolite Extraction

To extract metabolites from tissue slices, 700 μL of cold methanol was added to each tube, and samples were homogenized using the Retsch MM400 tissue homogenizer. 250 μL of the homogenate in methanol was transferred to a tube containing 100 uL of H_2_O with 100 ng Norvaline and 250 uL with 1 ug palmitate-d31. The aqueous and organic phases were collected and dried for analysis of polar metabolites and saponified fatty acids by GC-MS as previously described ^45^. The remaining interphase material was washed twice with cold MeOH and dried under air. 70 μL of the remaining homogenate was collected for protein quantification by BCA assay using the Pierce BCA protein assay kit (Pierce BCA protein assay kit, ThermoFisher Scientific, 23225). To extract neutral and polar lipids for liquid chromatography-mass spectrometry (LC-MS/MS), a deuterated internal standard mix containing EquiSPLASH (equivalent to 2 ng/sample per lipid), 50 pmol/sample of C15 Glucosyl(β) Ceramide-d7(d18:1-d7/15:0), 50 pmol/sample of C15 Lactosyl(β) Ceramide-d7(d18:1-d7/15:0), 100 pmol/sample of C18 Ganglioside GM3(d18:1/18:1-d3), and 100 pmol/sample C16 Globotriaosylceramide-d9 Gb3 (d18:1/16:0-d9) was added to the remaining homogenate in each tube. Extraction material was transferred to a new tube, vortexed for 5 minutes, and then centrifuged at 21,000 x g and 4 ^o^C for 10 minutes. The supernatant was transferred to a new tube and dried under N_2_ for LC-MS/MS analysis. All extracted samples were stored at -80 ^o^C prior to analysis by GC-MS or LC-MS/MS.

To extract metabolites from 2D cell cultures, media was aspirated at the desired time point, and wells were washed once with 0.9% saline. For the quantification of labeling in in the biomass, 250 μL of cold MeOH and 100 μL of H_2_O containing 100 nmol/sample of Norvaline was added directly to the well. Cells were scraped into Eppendorf tubes, and 35 μL of the lysate was collected for BCA protein quantification. 250 μL of cold CHCl_3_ was added to each sample. The aqueous and organic phases were collected and dried for analysis of polar metabolites and saponified fatty acids by GC-MS as previously described ^45^, and the remaining interphase material was washed twice with cold MeOH and dried under air. For 2D studies requiring both GC-MS and LC-MS/MS analyses, 700 μL of cold MeOH was added directly to the well. Cells were scraped from the plate. The lysate was transferred to an Eppendorf tube, and the extraction proceeded according to the tissue slice protocol.

### GC-MS Analysis of Polar Metabolites and Saponified Fatty Acids

Dried aqueous samples were processed for the analysis of polar metabolites including TCA cycle intermediates and free amino acid pools as previously described ^45^. Briefly, samples were derivatized with 2% methoxyamine in pyridine and N-tert-butyldimethylsily-N-methyltrifluoroacetamide (MTBSTFA) with 1% tBDMS to form methoxime-*tert*-butyldimethylchlorosilane (TBDMS) derivatives. GC-MS analysis was performed on the derivatized samples using a DB-35MS column (30 m by 0.25 mm i.d. by 0.25 μm; Agilent J&W Scientific) installed in an Agilent 7890B GC system integrated with an Agilent 5977a MS.

Fatty acids within the dried organic phase were saponified and analyzed per previously published methods ^45,83^. Fatty acids were saponified into fatty acid methyl esters (FAMEs) by incubation with 500 μL of 2% H_2_SO_4_ in MeOH for 2 hours. The resulting FAMEs were extracted through the addition of 100 μL of saturated NaCl solution and 500 μL of hexane. The hexane phase was collected, and GC-MS analysis was performed using a Select FAME column installed in an Agilent 7890 A GC system integrated with an Agilent 5975 C MS. The relative abundance of fatty acids was determined through normalization to the palmitate-d31 internal standard and the protein concentration in the sample.

GC-MS data for polar metabolites and saponified fatty acids was corrected for natural isotope abundance, as previously described in Fernandez, et al, using in-house scripts ^88^.

### Acid Hydrolysis and GC-MS Analysis of Biomass

Acid hydrolysis of biomass to free amino acids, nucleobases, and sugars was performed as previously described ^89^. Briefly, 500 μL of 6N HCl was added to dried interphases, and samples were incubated at 80 ^o^C for 2 hours. Samples were dried under air, then resuspended in 200 μL of 8:1 methanol:H_2_O. 40 μL of each sample was transferred to a fresh vial and dried prior to GC-MS analysis.

For analysis, samples were derivatized with 2% methoxyamine in pyridine and N-methyl-N-(trimethylsilyl) trifluoroacetamide (MSTFA) to form methoxime-trimethylsilyl (TMS) derivatives as previously described ^89^. GC-MS analysis was performed on the derivatized samples using a DB-35MS column (30 m by 0.25 mm i.d. by 0.25 μm; Agilent J&W Scientific) installed in an Agilent 7890B GC system integrated with an Agilent 5977a MS. Oven parameters were as follows: 1 minute at 90 ^o^C, increase 3 ^o^C/min to 150 ^o^C, 1.5 ^o^C/min to 180 ^o^C, 3 ^o^C/min to 200 ^o^C, 15 ^o^C/min to 300, then hold for 4 minutes at 300 ^o^C. GC-MS data for hydrolyzed biomass was corrected for natural isotope abundance using in-house scripts.

Acid hydrolysis of the biomass will release monosaccharide units that can then be analyzed on GC-MS. As acid hydrolysis is non-specific, other labile groups may also be hydrolyzed, including the N-Acetyl groups on GlcNAc and GalNAc. Thus, the GlcN pool will represent both GlcNAc and GlcN residues on glycans within the biomass. While analysis of monosaccharides as TMS-oximes derivatives eliminates the need to quantify multiple anomeric and cyclic/acyclic peaks, syn- and antimethoxime isomers are formed ^90^. There is a strong preference towards one isomer for mannose (Man), galactose (Gal), glucose (Glc), and N-Acetylglucosamine (GlcNAc) (data not shown). For fucose (Fuc), only the preferred isomer was quantifiable at the abundances present in KPC tissue slices. As expected, the labeling of these analytically induced isomers was essentially identical (data not shown), so only the most abundant isomer for each sugar will be shown for simplicity.

### LC-MS Analysis of Nucleotide Sugars

Nucleotide sugars were quantified from the dried aqueous fraction prior to GC-MS polar analysis. Analysis was performed on a Vanquish Flex UHPLC via HILIC-HRMS using an InfinityLab Poroshell 120 HILIC-z column (2.1 x 100mm, 2.7 μm, Agilent). The column temperature was maintained at 45°C. Mobile phases consisted of 10 mM ammonium carbonate in water (A) and in 95:5 acetonitrile: water (B) with 5μM medronic acid. The flow rate was 0.3 mL/min. The gradient started at 100% B, decreasing to 90% B over 4 min, to 50% B over 6 min, then to 30% B in 0.5 min, and then was held for 1 min before returning to 100% B and equilibrating for 8 min. Data was acquired via a Q-Exactive orbitrap mass spectrometer, with HESI conditions and MS parameters as described in Kambhampati, et al. ^91^. Analysis was performed using El-MAVEN, and data were corrected for natural isotope abundance sing in-house scripts ^88^.

### LC-MS/MS Analysis of Complex Lipids

Polar lipid analysis, including glycerophospholipids and sphingolipids, was performed using a Kinetex C18 column, 100 × 2.1 mm, 1.7 μm particle (Phenomenex) column installed on a Q Exactive orbitrap mass spectrometer with a Vanquish Flex Binary UHPLC (ultra-high-performance liquid chromatography) system (Thermo Fisher Scientific) as previously described ^52^. Liquid chromatography was performed with a column temperature of 35 ^o^C using a gradient of 98:2 v/v water: methanol with 5 mM ammonium acetate (mobile phase A) and 50:50 v/v methanol: 2-propanol with 5 mM ammonium acetate (mobile phase B) at 0.2 ml/min. Gradient parameters were as follows: 0 min, 30%B; 1 min, 30%B; 2 min, 70%B; 11 min, 95%B; 17 min, 30%B; 21.5 min, 30%B; 27 min, 30%B. Lipids were analyzed in positive mode with spray voltage of 3.2 kV. Sweep gas flow was 1 arbitrary units, auxiliary gas flow 2 arbitrary units and sheath gas flow 40 arbitrary units, with a capillary temperature of 325 °C. Full mass spectrometry (scan range 200–2,000 m/z) was used at 140,000 resolution with 10E6 automatic gain control and a maximum injection time of 100 ms. Data dependent MS2 (Top 12) mode at 17,500 resolution with automatic gain control set at 10E5 with a maximum injection time of 50 ms was used.

Samples were redried and resuspended in 13:6:1 Acetonitrile:isopropanol:H_2_O for the analysis of neutral lipids. As previously described in Kuna, et al. ^92^, LC-MS/MS analysis was performed on the Q Exactive using an Accucore C30, 150 by 2.1 mm, 2.6-μm particle (Thermo Fisher Scientific) column. Chromatography was performed using a gradient of 40:60 (v/v) water:acetonitrile with 10 mM ammonium formate and 0.1% formic acid and 10:90 (v/v) acetonitrile: 2-propanol with 10 mM ammonium formate and 0.1% formic acid. Lipids were analyzed with positive mode ionization, and MS2 data were generated using Top 12 mode.

All lipid data were analyzed using El-MAVEN. Relative abundances were calculated through normalization to the EquiSPLASH internal standard mix or other internal standards and to total protein content. For LPC species, the more abundant SN1 isoform is presented in figures. For DGs and applicable TGs, acyl chain identities are determined through MS2 spectra and are presented in arbitrary order. Natural isotope abundance correction was performed using in-house scripts.

### scRNASeq

Single cell data were extracted from NCBI or Zenodo (GSE125588, GSE129455, CA001063, and GSE155698). Data were processed using the Seurat package (version 5.1.0) in R, with quality control measures to exclude cells expressing fewer than 50 or more than 4,000 ∼ 6,000 genes, or with mitochondrial gene content exceeding 15%. Data normalization was performed using the LogNormalize method, and batch effects were addressed using Harmony. Principal component analysis (PCA) was applied to identify significant dimensions, and uniform manifold approximation and projection (UMAP) was used for visualization. Then, clustering was performed using shared nearest neighbor modularity optimization at varying resolutions to delineate biologically relevant subpopulations. Marker genes were identified through differential expression analysis, and clusters were annotated using established cell-type markers. If the NCBI data is not a filtered count, doublets were identified and removed using DoubletFinder, with homotypic proportions and sample-specific formation rates (0.087) incorporated into adjustments. Visualizations, including dimensionality reduction plots and violin plots, were created using Seurat and ggplot2. An interactive app, ShinyCell, was utilized to facilitate exploration of gene expression patterns and cluster annotations, allowing dynamic interrogation of the dataset.

### Immunohistochemistry and Quantification

Primary tumor material or post-culture slices were fixed in 10% neutral buffered formalin and paraffin-embedded. H&E staining was performed by the UCSD Tissue Technology Shared Resource according to standard protocols. For other IHC, 5 μm sections were deparaffinized with xylene and rehydrated sequentially in ethanol. Antigen retrieval was performed using sodium citrate buffer. Slides were treated with 3% H_2_O_2_ in TBST for 15 minutes to block endogenous peroxidase activity then blocked with 5% normal goat serum in 2.5% horse serum for 1 hour at room temperature (RT). Slides were incubated in primary antibody overnight at 4 ^o^C and appropriate secondary antibody for 30 minutes at RT. Slides were stained with DAB reagent and hematoxylin, then dehydrated in ethanol and xylene prior to mounting for imaging.

QuPath v.0.5.1 was used for quantification of immunohistochemistry staining. Hematoxylin and DAB stain vectors were set using the estimate stain vectors function. Optical density sum was used for cell detection and DAB threshold was set to count positive nuclei for Ki67 staining. For podoplanin and F4/80 quantification, a pixel classifier was used to threshold positive DAB staining. The positive area was then divided by the total hematoxylin positive area to quantify % positivity.

### Western Blotting

Cells were lysed in RIPA buffer at 70-80% confluency following culture. Lysates were clarified by centrifugation at 20,000 x g for 10 min at 4°C. Protein was quantified by BCA assay. 40 μg of protein was prepared in RIPA buffer with NuPAGE LDS Sample Buffer and Bolt Sample Reducing Agent, then boiled for 5 min. Protein was run on 4-12% NuPAGE Bis-Tris gel and transferred to PVDF membrane using BioRad Mini Trans-Blot Cell for 20V overnight at 4°C. Membrane was blocked in 5% milk in TBS-T for 1 hr at room temperature and primary antibody was incubated in 5% BSA in TBS-T at 4°C overnight. Membrane was then incubated with IRDye 800CW goat anti-mouse or anti-rabbit IgG secondary antibody (Li-Cor #926-32210, #926-32211) for 1 hr at room temperature.

### Lipid Modeling

Isotopomer spectral analysis (ISA) to estimate the percentage of newly synthesized palmitate was performed using the INCA MFA software ^93^. The model was fit to experimentally-obtained and natural abundance corrected palmitate labeling data by simulation, assuming the synthesis of one palmitate molecule from eight acetyl-CoA molecules ^82,92^. Flux value estimation was performed on all replicates concurrently. 95% confidence intervals were calculated via parameter continuation, and statistical significance (p<0.05) was determined by a lack of overlap.

Lipid MFA to estimate the labeling of relevant precursor pools (i.e., palmitate, Neu5Ac), fatty acid elongation fluxes, and synthetic fluxes for LPCs, SMs, and GSLs was performed as described in Wessendorf-Rodriguez, et al. ^52^ Metabolic networks were defined for the synthesis of complex lipids of interest, and the model was fit to experimentally-obtained and natural abundance corrected isotopologue data for key lipids and intermediates. Analyses assumed metabolic steady state, and precursors were assumed to be at isotopic steady state or pseudo-steady state with respect to product labeling. These assumptions were supported by metabolite data within the timeframe investigated. For estimates of elongation fluxes, the metabolic network included LPC and SM species containing 16:0 and 24:0 acyl chains: LPC(16:0), LPC(24:0), SM(34:1), and SM(42:1), and included labeling data from relevant DHCer and Cer intermediates as well as labeling for the Glyc3P and Ser pools. Flux value estimation and sensitivity analysis for determining 95% confidence intervals was performed on all replicates within a condition concurrently. For GSLs, the metabolic network contained GM3(d18:1/16:0) and Gb4(d18:1/16:0), with labeling data from DHCer(d18:1/16:0), Cer(d18:1/16:0), and LacCer(18:1/16:0).

