## Supplemental Figures for "Metabolic Salvage and Acyl-chain Remodeling Support Glycosphingolipid Synthesis within the PDAC Tumor Microenvironment"

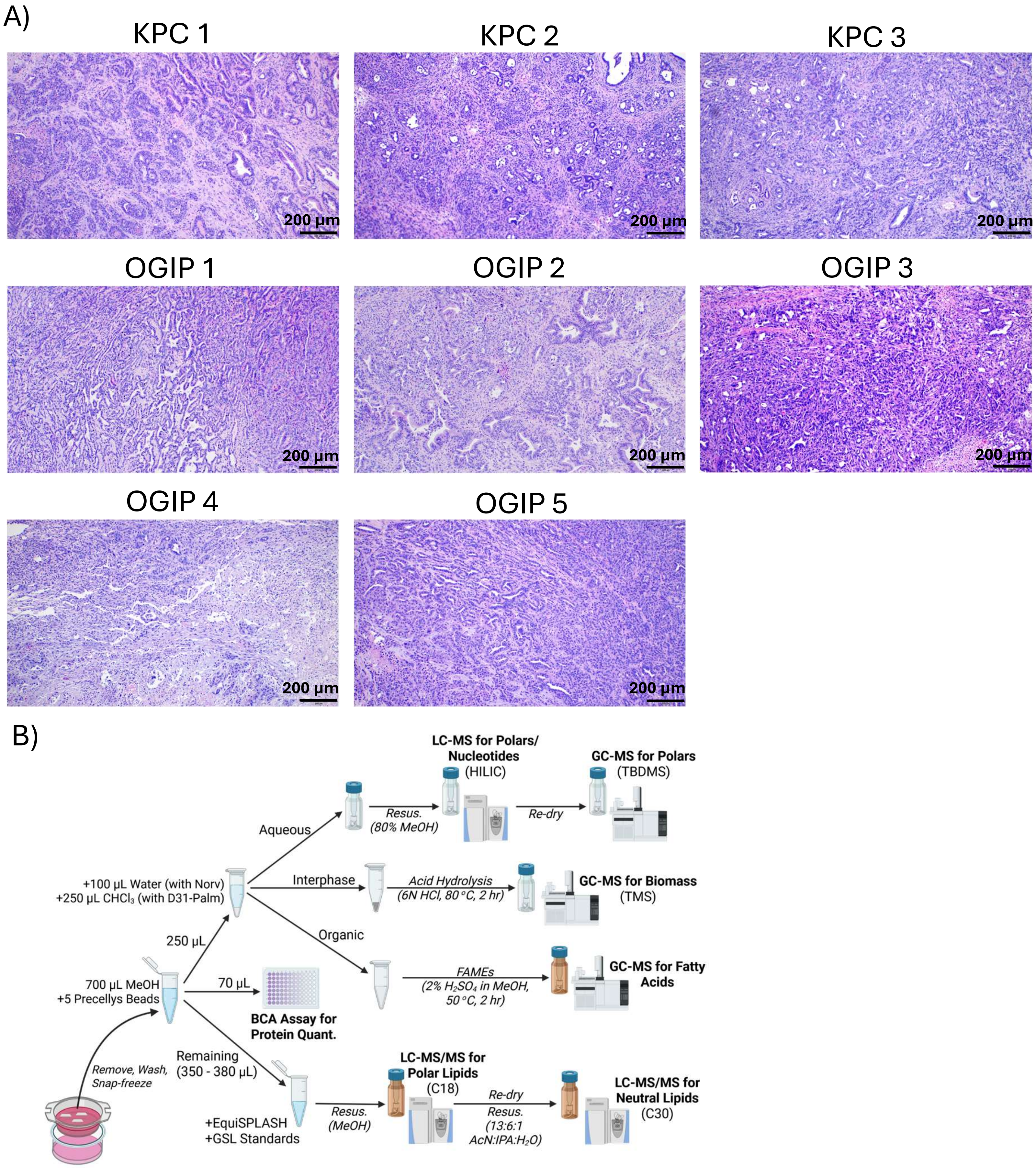


**Extended Data Fig. 1. KPC and KPC OGIP Tumors in *Ex Vivo* Culture** (A) Representative H&E images from KPC and KPC OGIP tumors used in the subsequent studies. Scale bars show 200 μm. (B) Schematic describing the partitioning of tissue slice homogenate for multiple analyses, including extraction solvents and column chemistry or derivatization, as relevant. (GSL, Glycosphingolipid; IS, Internal Standard; Norv, Norvaline; Palm, Palmitate)

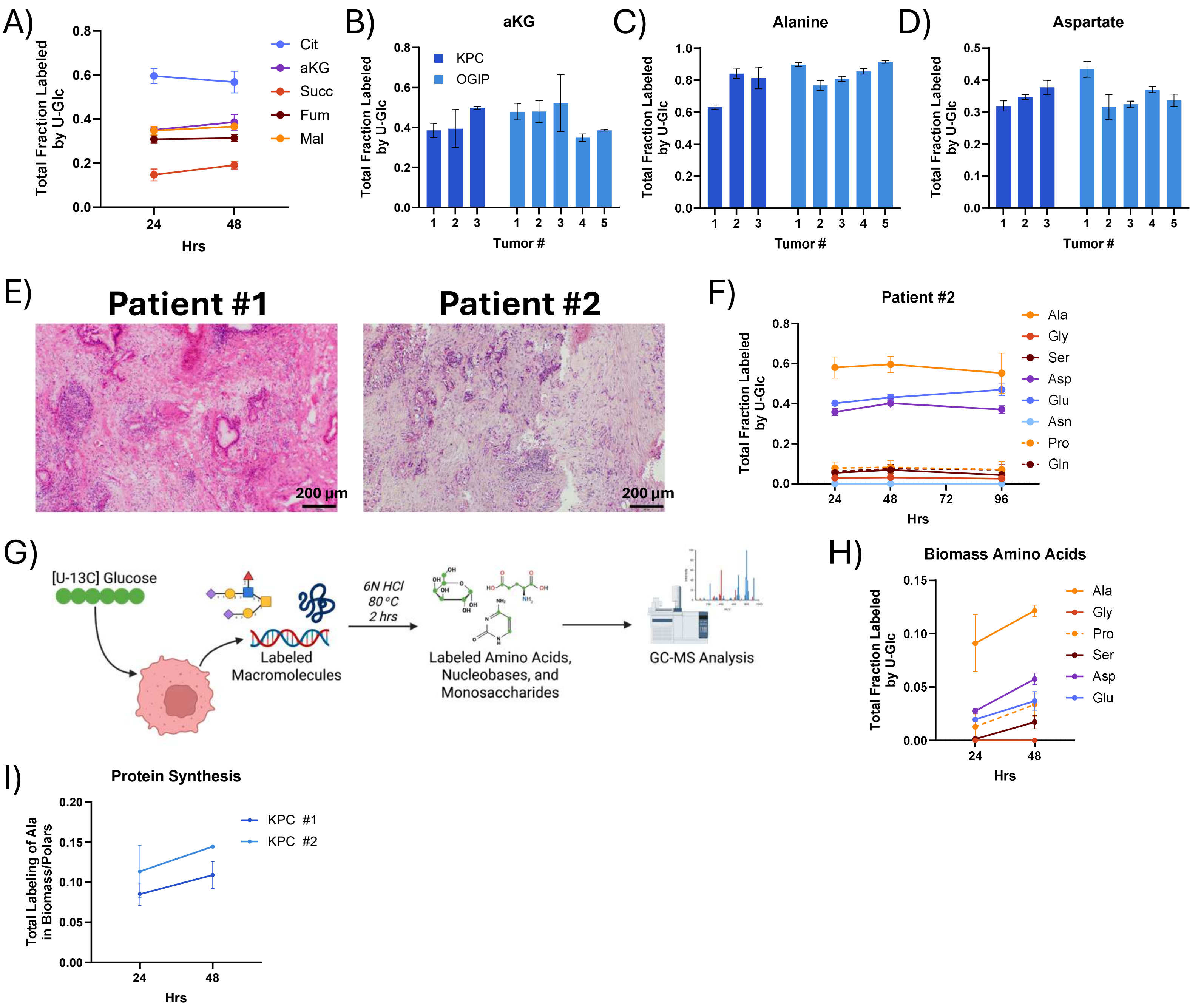


**Extended Data Fig. 2. Protein Synthesis in PDAC Tissue Slices.** (A) Total fraction labeled of TCA cycle intermediates from U-Glc in tumor slices from a representative KPC tumor (KPC #2). Total fraction of (B) alpha-ketoglutarate (aKG), (C) alanine, and (D) aspartate labeled by U-Glc at 48 hours in tumor slices. (E) H&E images for human PDAC tumors used in metabolic analysis. Scale bars represent 200 µm. (F) Non-essential amino acid labeling from U-Glc in tissue slices from Patient #2. (G) Schematic demonstrating how stable isotope tracing of hydrolyzed biomass can be used to quantify macromolecule synthesis. (H) Total fraction labeled by U-Glc of non-essential amino acids in the hydrolyzed biomass of tissue slices from KPC #2. (I) Protein synthesis over time in KPC tumor slices calculated as the total enrichment of alanine labeled in the hydrolyzed biomass normalized to the total enrichment of the polar alanine pool.

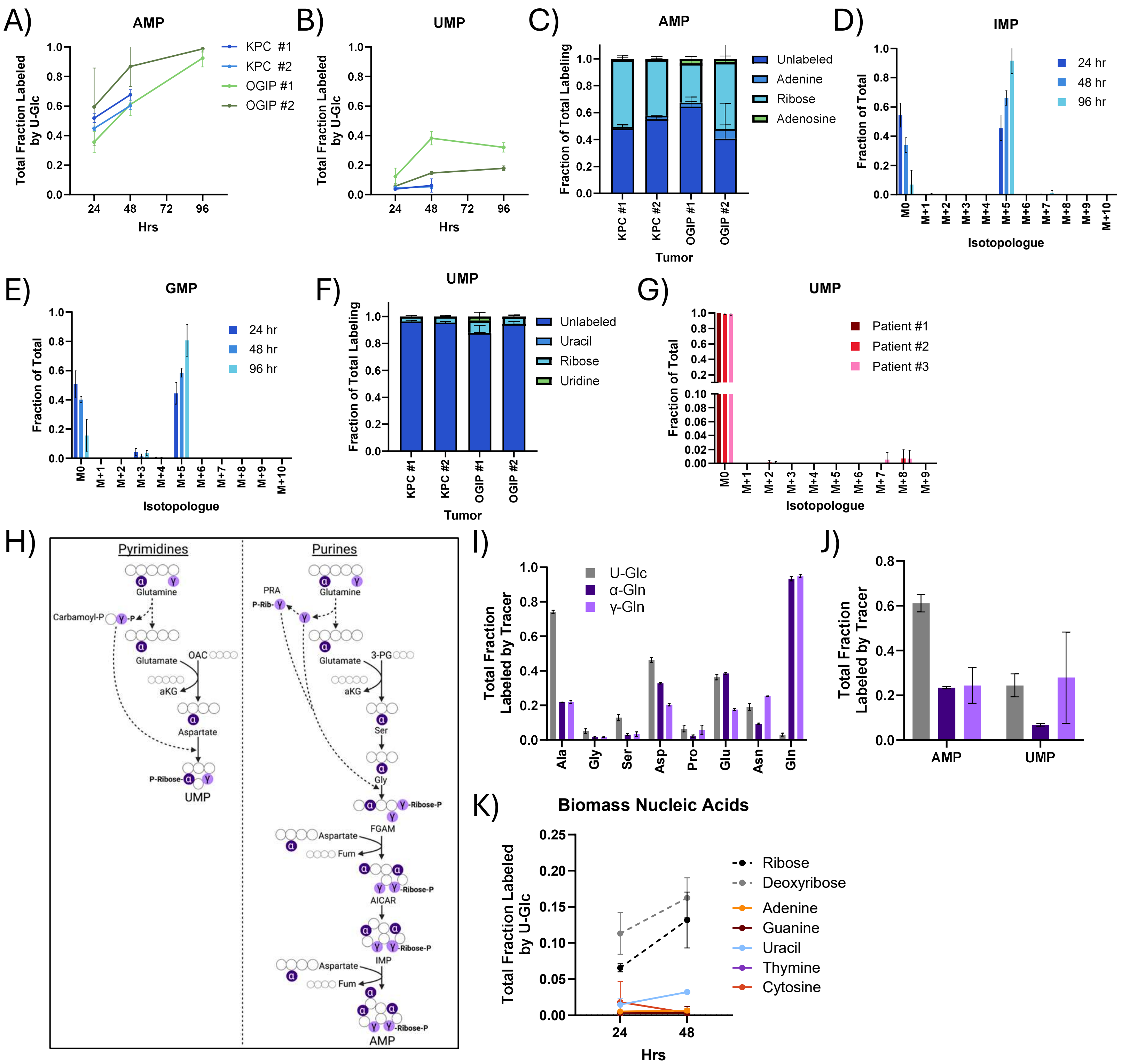


**Extended Data Fig. 3. Nucleotide and Nucleic Acid Synthesis in PDAC Tumor Slices.** Total labeling of (A) AMP and (B) UMP from U-Glc in tumor slices. (C) Summary of AMP isotopologue distribution from U-Glc at 24 hours. (D) IMP and (E) GMP isotopologue distributions from U-Glc in tissue slices from KPC OGIP #2. (F) Summary of the isotopologue distributions of UMP from U-Glc at 24 hours. (G) Isotopologue distribution of UMP from 96 hours of U-Glc tracing in human PDAC slices. (H) Diagram showing the incorporation of nitrogen from L-glutamine into UMP and AMP synthesis. (I) Total labeling of non-essential amino acids from U-Glc or $^{15}$N Glutamine tracers at 48 hours in tissue slices from a KPC tumor. (J) Total labeling of AMP and UMP via tracing with U-Glc or $^{15}$N-Glutamine for 48 hours in tissue slices from a KPC tumor. (K) Total labeling of ribose, deoxyribose, and nucleobases in the hydrolyzed biomass by U-Glc in tissue slices from KPC #2.

All means represented three slices from a single tumor. All error bars show standard deviation.

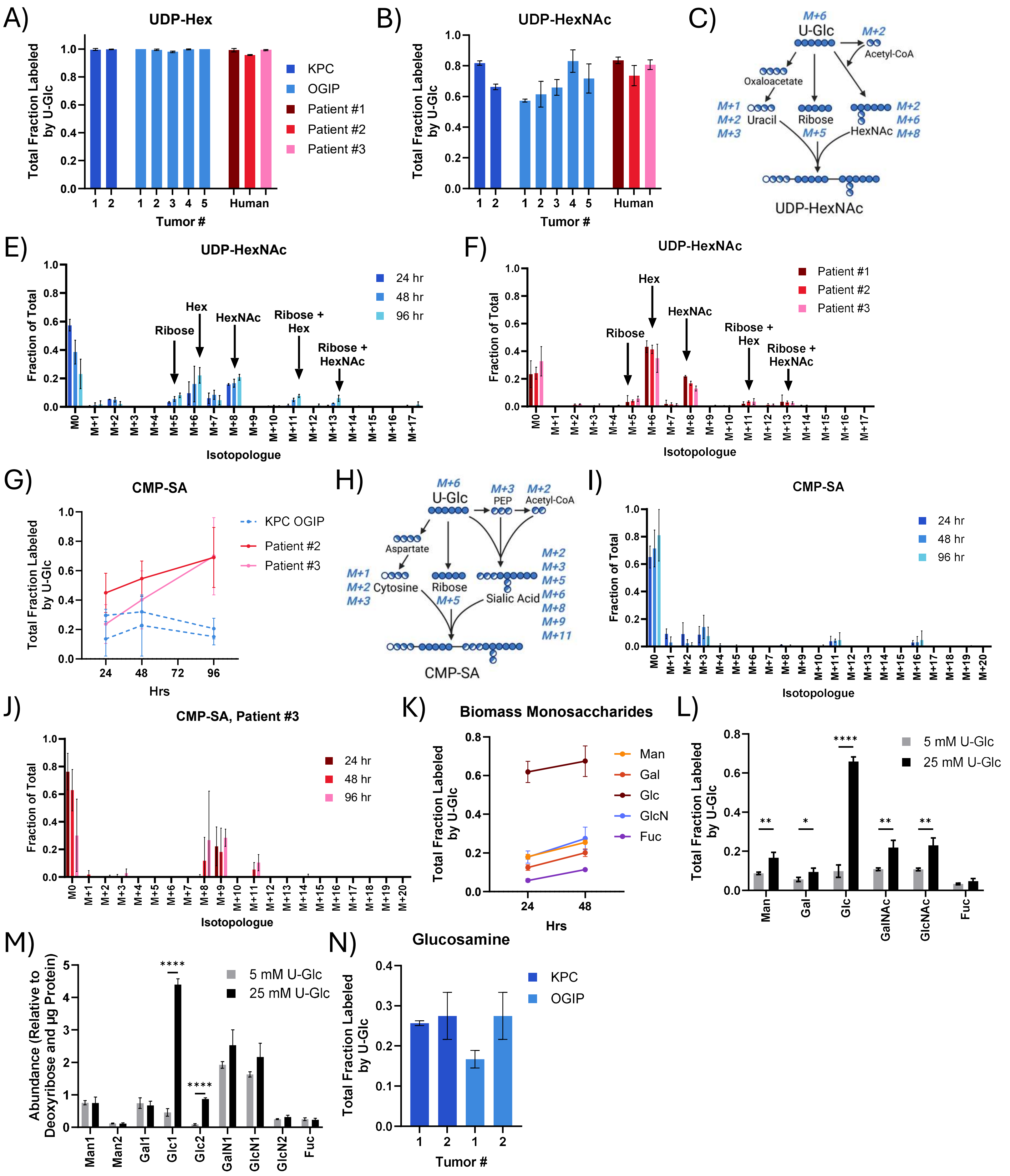
A) UDP-Hex
Total Fraction Labeled by U-Glc
Tumor #
Human
KPC
OGIP
Patient #1
Patient #2
Patient #3
B) UDP-HexNAc
C) M+6 U-Glc
M+2 Acetyl-CoA
Oxaloacetate
M+1 M+2 M+3 Uracil
Ribose M+5
HexNAc M+2 M+6 M+8
UDP-HexNAc
E) UDP-HexNAc
Fraction of Total
24 hr
48 hr
96 hr
Ribose
Hex
HexNAc
Ribose + Hex
Ribose + HexNAc
Isotopologue
F) UDP-HexNAc
G) CMP-SA
KPC OGIP
Hrs
H) M+6 U-Glc
M+3 PEP
M+2 Acetyl-CoA
Aspartate
M+1 M+2 M+3 Cytosine
Ribose M+5
Sialic Acid
M+2 M+3 M+5 M+6 M+8 M+9 M+11
CMP-SA
I) CMP-SA
J) CMP-SA, Patient #3
K) Biomass Monosaccharides
Man
Gal
Glc
GlcN
Fuc
L) 5 mM U-Glc
25 mM U-Glc
GalNAc
GlcNAc
M) Abundance (Relative to Deoxyribose and µg Protein)
Man1
Man2
Gal1
Glc1
Glc2
GalN1
GlcN1
GlcN2
N) Glucosamine

**Extended Data Fig. 4. Glycan Synthesis in PDAC Tumor Slices.** Total labeling of (A) UDP-Hexose (UDP-Hex) and (B) UDP-N-acetylhexosamine (UDP-HexNAc; UDP-GlcNAc and UDP-GalNAc). (C) Diagram demonstrating the possible labeling of UDP-HexNAc from U-Glc tracing. Isotopologue distribution of UDP-HexNAc from U-Glc in (E) tissue slices from KPC OGIP #2 and (F) human PDAC slices at 24 hours. (G) Total CMP-SA labeling from U-Glc in tissue slices. CMP-SA was not quantifiable in slices from Patient #1. (H) Diagram demonstrating the possible labeling of CMP-SA from U-Glc tracing. Isotopologue distribution of CMP-SA from U-Glc in tissue slices from (I) KPC OGIP #2 and (J) Patient #3. (K) Total labeling of sugars in the hydrolyzed biomass by U-Glc in tissue slices from KPC #2. (L) Total labeling and (M) abundance of sugars in the hydrolyzed biomass of KPC tissue slices cultured at different concentration of U-Glc for 24 hours. Monosaccharide abundances are normalized to the abundance of deoxyribose and total protein in the sample. (N) Total labeling of glucosamine in hydrolyzed biomass glycans from KPC tumor slices. Hydrolyzed glucosamine pool includes N-acetyl-glucosamine.

All means represented three slices from a single tumor. All error bars show standard deviation. Where relevant, statistical significance was determined by two-tailed *t*-test (*, $P < 0.05$; **, $P < 0.01$; ***, $P < 0.001$; ****, $P < 0.0001$).

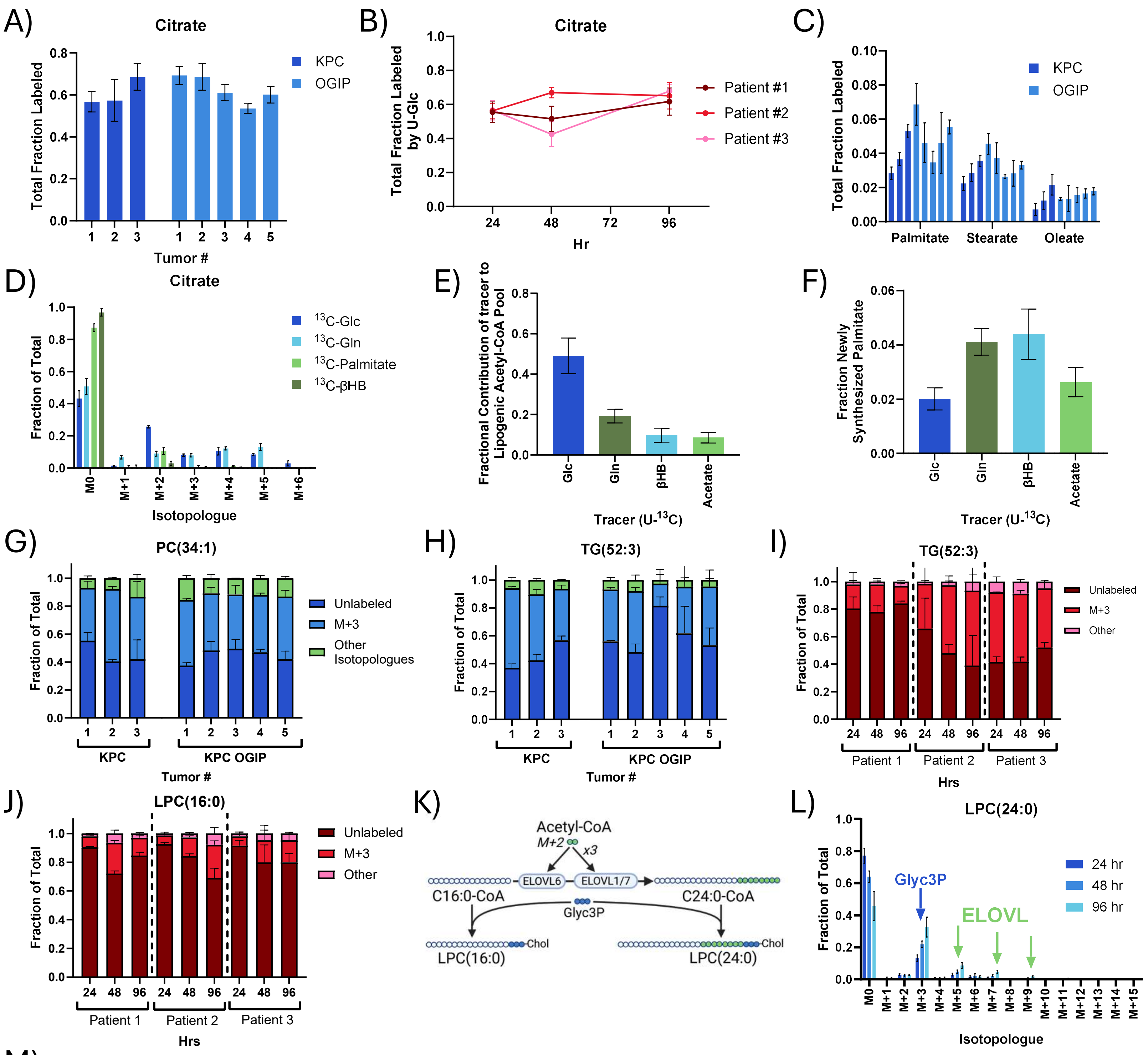

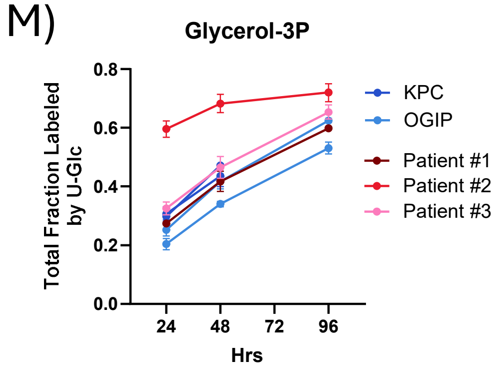


**Extended Data Fig. 5. Glycerolipid Synthesis in PDAC Tumor Slices.** (A) Total fraction of citrate labeled by U-Glc at 48 hours in KPC tumor slices. (B) Total citrate labeling in human PDAC tumor slices. (C) Total labeling of the palmitate, stearate, and oleate in the saponified lipid fraction by U-Glc at 48 hours in tumor slices. (D) Isotopologue distribution for citrate from $^{13}C$ labeled lipogenic carbon sources at 48 hours in tissue slices from KPC #1. (E) Fractional contribution of $^{13}C$ labeled tracers to the lipogenic acetyl-CoA pool at 48 hours and (F) palmitate synthesis from $^{13}C$ labeled lipogenic carbon sources at 48 hours, as calculated via isotopomer spectral analysis from the saponified palmitate pool. Error bars represent 95% confidence intervals via parameter continuation. (G) PC(34:1) and (H) TG(52:3) isotopologue distributions from U-Glc tracing in KPC tumor slices at 48 hours. (I) TG(52:3) and (J) LPC(16:0) isotopologue distributions from U-Glc in human PDAC tumor slices over time. (K) Diagram depicting the elongation of fatty acids by ELOVL elongases and their subsequent incorporation into LPCs. (L) Isotopologue distribution of LPC(24:0) from U-Glc in KPC OGIP #2 tissue slices. Isotopologues likely derived from the incorporation of labeled Glyc3P or via fatty acid elongations (ELOVL) are highlighted. (M) Total labeling of Glyc3P from U-Glc in PDAC tumor slices over time.
All means represented three slices from a single tumor. Unless otherwise noted, all error bars show standard deviation.

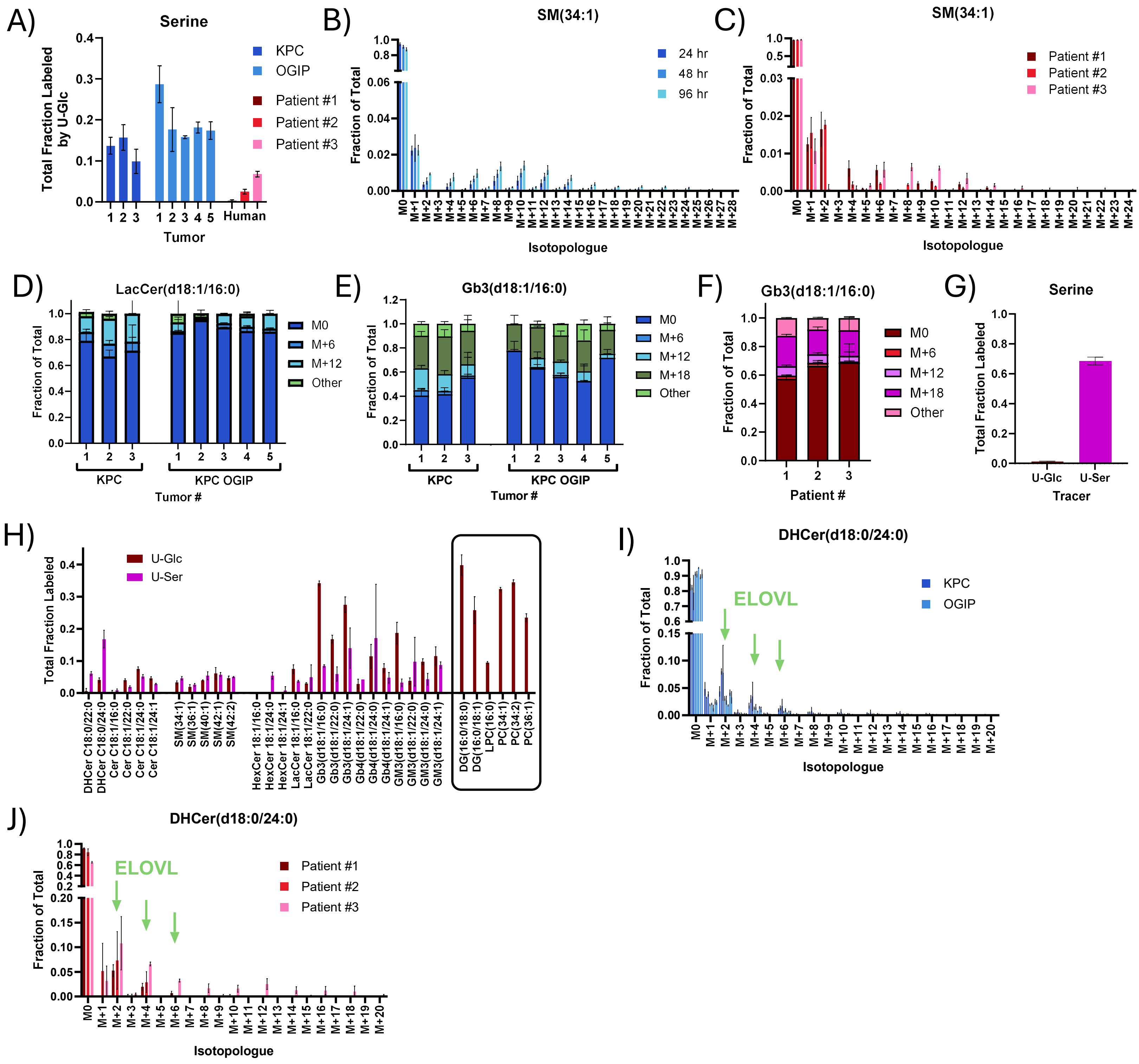


**Extended Data Fig. 6. Sphingolipid Synthesis in PDAC Tumor Slices**. (A) Total labeling of Serine from U-Glc at 48 hours in PDAC tumor slices. Isotopologue distribution from U-Glc for sphingomyelin(34:1) in (B) KPC OGIP #2 and (C) in human PDAC slices at 48 hours. Isotopologue distribution of (D) LacCer(d18:1/16:0) and (E) Gb3(d18:1/16:0) from U-Glc in KPC tumor slices at 48 hours. (F) Isotopologue distribution for Gb3(d18:1/16:0) in human PDAC slices at 48 hours. (G) Total labeling of serine from uniformly labeled $^{13}C$ tracers at 24 hours in tissue slices from Human Patient #1. (H) Total labeling of key sphingolipids from 24 hours of tracing with U-Glc or U-Ser in tumor slices from Patient #1. U-Glc labeling of key glycerolipid species at 24 hours is shown for comparison. (I) DHCer(d18:0/24:0) isotopologue distribution from U-Glc tracing in tumor slices and (J) in human PDAC tissue slices at 48 hours.
All means represented three slices from a single tumor. All error bars show standard deviation.

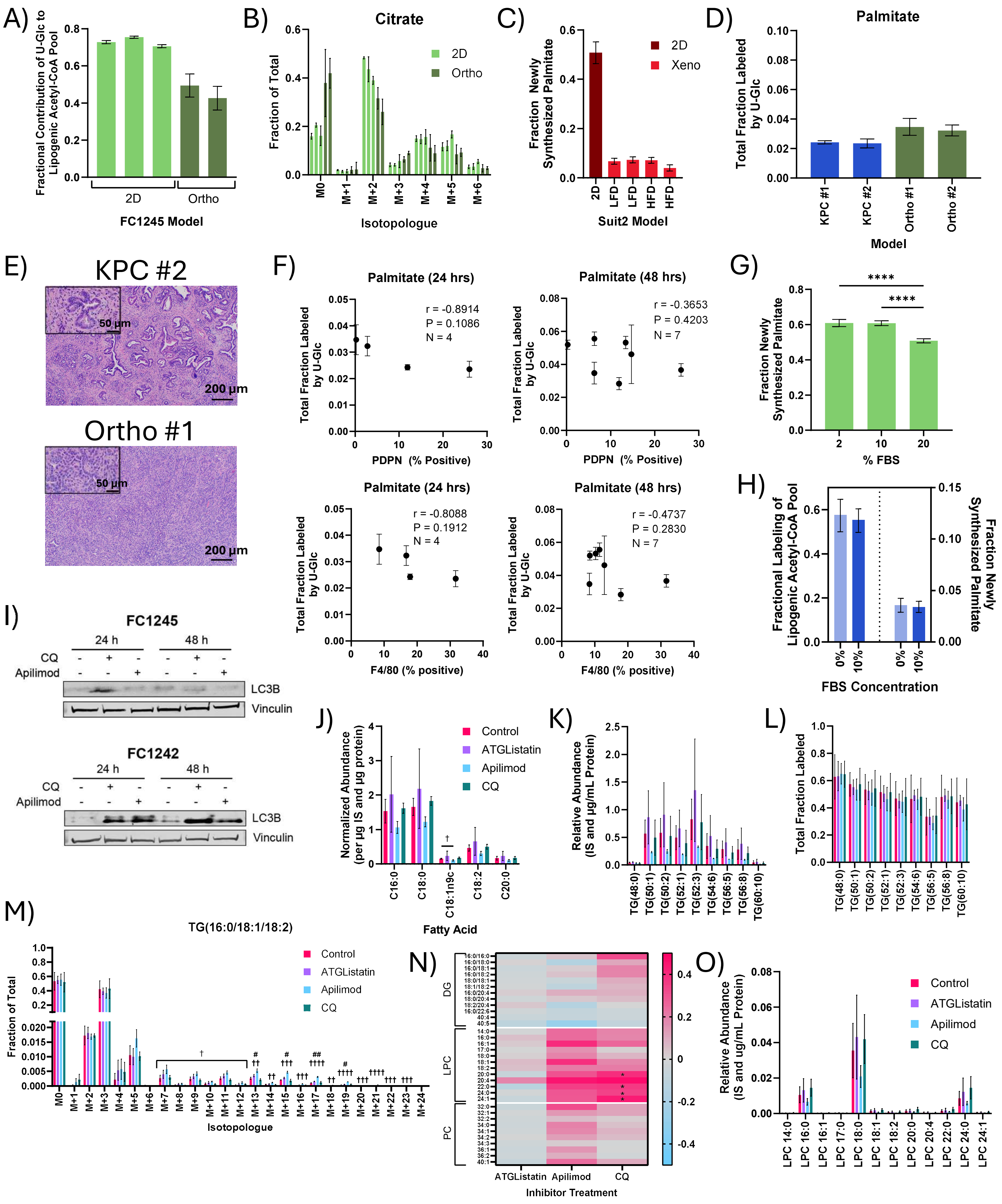

A)
Fractional Contribution of U-Glc to Lipogenic Acetyl-CoA Pool
2D
Ortho
FC1245 Model
B)
Citrate
Fraction of Total
2D
Ortho
M0
M+1
M+2
M+3
M+4
M+5
M+6
Isotopologue
C)
Fraction Newly Synthesized Palmitate
2D
Xeno
2D
LFD
LFD
HFD
HFD
Suit2 Model
D)
Palmitate
Total Fraction Labeled by U-Glc
KPC #1
KPC #2
Ortho #1
Ortho #2
Model
E)
KPC #2
50 µm
200 µm
Ortho #1
50 µm
200 µm
F)
Palmitate (24 hrs)
Total Fraction Labeled by U-Glc
r = -0.8914
P = 0.1086
N = 4
PDPN (% Positive)
Palmitate (48 hrs)
r = -0.3653
P = 0.4203
N = 7
PDPN (% Positive)
Palmitate (24 hrs)
r = -0.8088
P = 0.1912
N = 4
F4/80 (% positive)
Palmitate (48 hrs)
r = -0.4737
P = 0.2830
N = 7
F4/80 (% positive)
G)
Fraction Newly Synthesized Palmitate
****
****
% FBS
H)
Fractional Labeling of Lipogenic Acetyl-CoA Pool
Fraction Newly Synthesized Palmitate
0%
10%
0%
10%
FBS Concentration
I)
FC1245
24 h
48 h
CQ
Apilimod
LC3B
Vinculin
FC1242
24 h
48 h
CQ
Apilimod
LC3B
Vinculin
J)
Normalized Abundance (per µg IS and µg protein)
Control
ATGListatin
Apilimod
CQ
C16:0
C18:0
C18:1n9c
C18:2
C20:0
Fatty Acid
K)
Relative Abundance (IS and µg/mL Protein)
TG(48:0)
TG(50:1)
TG(50:2)
TG(52:1)
TG(52:3)
TG(54:6)
TG(56:5)
TG(56:8)
TG(60:10)
L)
Total Fraction Labeled
TG(48:0)
TG(50:1)
TG(50:2)
TG(52:1)
TG(52:3)
TG(54:6)
TG(56:5)
TG(56:8)
TG(60:10)
M)
TG(16:0/18:1/18:2)
Fraction of Total
Control
ATGListatin
Apilimod
CQ
Isotopologue
N)
DG
LPC
PC
ATGListatin
Apilimod
CQ
Inhibitor Treatment
O)
Relative Abundance (IS and ug/mL Protein)
Control
ATGListatin
Apilimod
CQ
LPC 14:0
LPC 16:0
LPC 16:1
LPC 17:0
LPC 18:0
LPC 18:1
LPC 18:2
LPC 20:0
LPC 20:4
LPC 22:0
LPC 24:0
LPC 24:1

**Extended Data Fig. 7. The presence of the tumor microenvironment influences PDAC lipid metabolism.** (A) Fractional contribution of U-Glc to the lipogenic acetyl-CoA pool at 48 hours in 2D cultures of FC1245 cells or in tissue slices from FC1245 orthotopic tumors (Orthos), as calculated via ISA of the saponified palmitate pool. (B) Isotopologue distribution for citrate in 2D FC1245 cells or in tissue slices of FC1245 orthotopic tumors. (C) Palmitate synthesis by ISA modeling of saponified palmitate for Suit2 cells in 2D culture vs orthotopic xenografts (Xeno). Increased dietary fat consumption may decrease palmitate synthesis, but synthesis remains much lower than 2D cell culture (LFD, low fat diet; HFD, high fat diet). (D) Total fraction of palmitate labeled by U-Glc at 24 hours in KPC tumors and in FC1242 orthotopic tumors shown in Fig 6C-D. (E) Representative H&E images of primary tumor material from KPC #2 and FC1242 Ortho #1. Scale bars show 200 µm. Inset scale bars are 50 µm. (F) Pearson correlations for percentage podoplanin- (PDPN) or F4/80-positive cells and total palmitate labeling from U-Glc at 24 and 48 hours. (G) Palmitate synthesis via ISA modeling of the saponified palmitate pool after 48 hours of U-Glc tracing in 2D FC1245 cell cultures using varying concentrations of fetal bovine serum (FBS). (H) Fractional contribution of U-Glc to the lipogenic acetyl-CoA pool (left) and palmitate synthesis (right) at 48 hours from the saponified pamitate pool via ISA in tissue slices from KPC #1. (I) Western blot for autophagy-associated protein LC3B in 2D PDAC cell lines treated with 25 µM CQ or 100 nM apilimod. (J) Relative abundance of fatty acids in the saponified lipids after 48 hours of culture with 10 µM ATGListatin, 100 nM apilimod, or 25 µM CQ in tissue slices from KPC OGIP #5. (K) Relative abundance and (L) total fraction labeled of key TG species at 48 hours with inhibitor treatments. (M) Isotopologue distribution for TG(16:0/18:1/18:2) (TG(52:3)) from 48 hours of U-Glc tracing. Acyl chains are verified by MS2 and are listed in arbitrary order. (N) Relative difference in the labeling of key glycerophospholipid species by U-Glc in KPC OGIP #5 tumor slices with 48 hours of inhibitor treatment. (O) Relative abundance of key LPC species at 48 hours.

All means represented three slices from a single tumor or three wells of 2D cell culture. For ISA results (Panels A, C, G, and H), all error bars represent 95% confidence intervals, and statistic significance is determined by overlapping error bars. For all other panels, error bars show standard deviation. Significance denoted as # for ATGListatin, † for Apilimod, * for CQ. (*, $P < 0.05$; **, $P < 0.01$; ***, $P < 0.001$; ****, $P < 0.0001$).

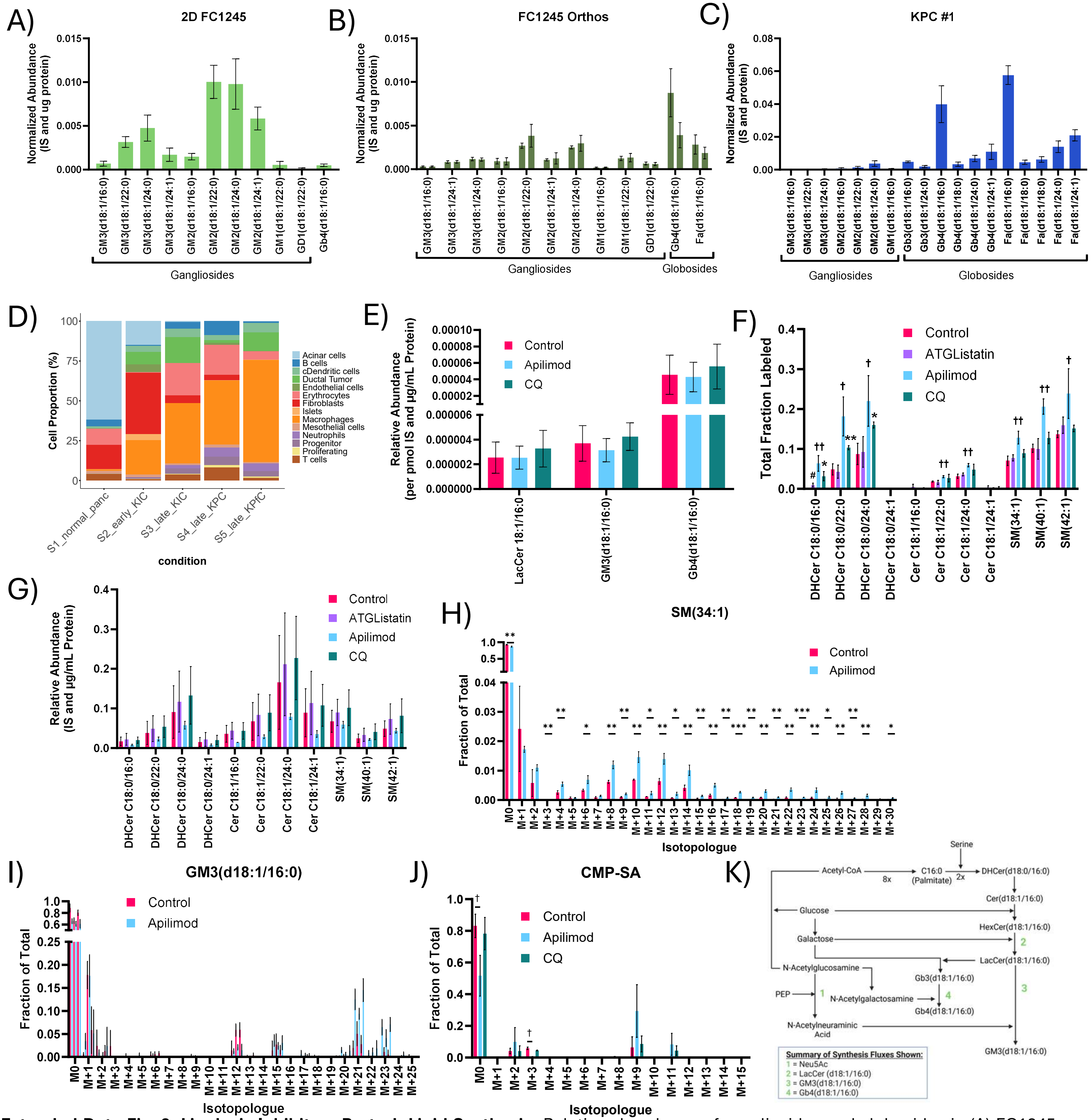


**Extended Data Fig. 8. Lipolysis Inhibitors Perturb Lipid Synthesis.** Relative abundance of gangliosides and globosides in (A) FC1245 cells in 2D culture, (B) tissue slices from FC1245 orthotopic tumors, and (C) tissues slices from KPC #1 after 24 hours of culture. Only species which could be quantified and MS2 verified are shown. (D) Proportions of each cell type represented in the scRNASeq dataset (GSE125588). (E) Relative abundance of key glycosphingolipids after 48 hours of treatment with 100 nM apilimod or 25 µM CQ. (F) Total fraction labeled and (G) abundance of key dihydroceramide (DHCer), ceramide (Cer), and sphingomyelin (SM) species from 48 hours of U-Glc tracing in tissue slices from KPC OGIP #5. (H) SM(34:1) isotopologue distribution for U-Glc tracing with apilimod treatment. Significance is denoted by *. (K) Isotopologue distributions of GM3(D18:1/16:0) from 48 hour of U-Glc tracing in tissue slices from three KPC OGIP tumors (#3, #4, and #5). Apilimod treated slices are shown immediately to the right of their respective untreated control slices. (J) Isotopologue distribution for CMP-SA from U-Glc after 48 hours of inhibitor treatment in tissue slices from KPC OGIP #5. (K) Diagram summarizing the network and key fluxes quantified using lipid MFA modeling for glycosphingolipid synthesis.

All means represented three slices from a single tumor, and all error bars represent standard deviation. Statistical significance was determined by two-tailed *t*-test. Significance denoted as # for ATGListatin, † for Apilimod, * for CQ (*, $P < 0.05$; **, $P < 0.01$; ***, $P < 0.001$; ****, $P < 0.0001$).
